# Structure of core fungal endobiome in *Ulmus minor*: patterns within the tree and across genotypes differing in tolerance to Dutch elm disease

**DOI:** 10.1101/2020.06.23.166454

**Authors:** David Macaya-Sanz, Johanna Witzell, Carmen Collada, Luis Gil, Juan Antonio Martin

**Affiliations:** Departamento de Sistemas y Recursos Naturales, ETSI Montes, Forestal y del Medio Natural, Universidad Politécnica de Madrid, Madrid, 28040 Spain; Department of Biology, West Virginia University, Morgantown, WV, 26506 USA; Faculty of Forest Sciences, Southern Swedish Forest Research Centre, Swedish University of Agricultural Sciences, Alnarp, Sweden

**Author notes:** Corresponding author: Juan Antonio Martin.

**Keywords:** fungal endophytes, metabarcoding, plant-fungal interactions, Dutch elm disease, core microbiome, tree microbiome

## Abstract

Plants harbour a diverse fungal community with complex symbiotic interactions and significant roles in host physiology. However, the cues that steer the composition and structure of this community are poorly understood. Trees are useful models for assessing these factors because their large size and long lifespan give these ecosystems time and space to evolve and mature. Investigation of well-characterised pathosystems such as Dutch elm disease (DED) can reveal links between endomycobiome and pathogens. We examined the endophytic mycobiome across the aerial part of a landmark elm tree to identify structural patterns within plant hosts, highlighting not only commonalities but also the effect of local infections in some branches of the crown. We used a common garden trial of trees with varying levels of genotypic susceptibility to DED to identify associations between susceptibility and endomycobiome. Three families of yeasts were linked to higher DED tolerance: Buckleyzymaceae, Herpotrichiellaceae and Tremellaceae. Surveying a natural population with a gradient of vitality, we found some taxa enriched in declining trees. By combining all surveys and adding a further study in a distant natural population, we found evidence of a *U. minor* core mycobiome, pervasive within the tree and ubiquitous across locations, genotypes and health status.

## Introduction

The endophytic community in annual plants and deciduous leaves is largely configured each season at early developmental stages, when priority effects in initial assemblages appear to be crucial [1]. However, in long-lived forest trees, interactions among fungi that invade perennial organs are likely to be more complex [2]. Studies in crop plants have reported consistent co-occurrence of endophytic assemblages, suggesting the existence of a core microbiome, i.e., a minimum composition of microbes that constantly reside in the plant and are shared among conspecific hosts [3]. This core microbiome appears to be structured in functional networks that can positively or negatively affect host performance [4]. However, little is understood about the extent to which the endophytic flora of perennial organs is defined by random colonisation processes or shaped by environmental cues or active host recruiting factors (creating a core microbiome as a result of plant-microbe coevolution) [5]. Because of their large size and long life, forest trees are suitable organisms for answering these questions and unravelling the taxonomic diversity and structure within and among hosts. However, perhaps because sampling in large tree crowns presents obvious methodological difficulties, the diversity and within-tree distribution of endophytes in long-lived trees remain largely unexplored.

Tree architecture (e.g. crown height, age structure of twigs, light availability and microclimate within the crown) appears to influence endophyte abundance [6, 7]. The endophytic mycobiome can vary with geographical location and host vitality [8]. Local climate conditions (e.g. temperature and rainfall) can strongly influence the prevailing fungal inoculum through environmental filtering [9, 10]. The composition and age of the surrounding vegetation and historical changes in land uses can also affect plant microbiomes. Some endophytes that colonise long-lived trees are facultative (or opportunistic) saprotrophs and necrotrophs living in a cryptic phase under healthy host development [11, 12]. Stress affecting the host physiological status could induce shifts in endophytic communities.

Tree resistance to disease is affected by multiple factors, including the genetic make-up of hosts, the virulence of pathogens, and the environment. Evidence increasingly suggests that endophytic infections trigger systemic responses in plants [13]. Endophytes may also modulate the outcome of host interactions with pathogens by producing antimicrobial compounds or competing with other microorganisms, leading to higher host tolerance to pathogens [14, 15]. Increasing evidence suggests that the core microbiome of a plant exerts key effects on plant performance and tolerance to various stressors [3, 16].

The current pandemic of Dutch elm disease (DED) is caused by the highly virulent pathogen *Ophiostoma novo-ulmi* Brasier. DED has caused massive loss of elm trees in Europe and North America. The disease is transmitted by elm bark beetles of the genera *Scolytus* and *Hylurgopinus*, or through root grafts. The fungus becomes established in internal plant tissues reaching the vascular system, where it spreads systemically, causing massive occlusion and cavitation of xylem vessels, ultimately leading to a wilt syndrome [17]. The diversity of endophytic fungi in elms remains largely unexplored. A previous study showed that endophyte frequency and diversity in elms were influenced by host location and genotype [18], and that the diversity of the fungal endobiome in xylem (but not in leaf or bark) tissues of clones susceptible to DED was higher than in tolerant clones. However, this study addressed only the culturable fraction of endophyte fungi, which was calculable in less than 5% of the total fungal richness within a tree (authors, personal observation).

We applied molecular, bioinformatic and statistical tools to appraise the structure of the endophytic mycobiome within and among individuals in *Ulmus minor*, a European elm species highly susceptible to DED, to determine the portion of mycobiome considered to be core. Using the outcomes of this analysis, we addressed four key questions: **i) Does endophyte diversity vary within the aerial part of the tree?** We studied the fungal endobiome of stem tissue samples distributed throughout the aerial part of a large *U. minor* tree more than 200 years old. **ii) Is the level of susceptibility to *O. novo-ulmi* linked to variation in endophyte composition?** We examined the endophyte diversity of 10 healthy *U. minor* trees with varying degrees of genotypic tolerance to DED. **iii) How is the fungal endobiome altered as tree vitality diminishes?** We explored the fungal composition of six large *U. minor* trees showing different vitality levels but growing in the same location. **iv) To what extent is the twig endophytic flora of elms affected by their growing location?** The local environment is known to exert a strong influence on endophyte composition. However, ubiquitous taxa associated with healthy elms could have important functions in plant performance, following the concept of core microbiome. The study includes elms located in four different sites in central Spain.

## Materials and Methods

### Plant material

To determine how tree stem fungal microbiome is structured, we sampled wood tissue from twigs (1-2 cm diameter) and trunks (5-cm cores at breast height) from trees at four locations in Spain. We focused on stem endobiome because it is a perennial tissue, in which microbiome interactions have time to evolve and mature, and because the agent responsible for DED is a vascular pathogen and therefore mostly interacts with the xylem microbiome. To prevent inclusion of epiphytic flora, the external layer of the bark (periderm) was manually extirpated. The stem tissues analysed were xylem and the remaining phloem.

#### i) Within-tree mycobiome variation

Ten locations were sampled in a landmark *Ulmus minor* tree (Somontes, Madrid, Spain; Fig. 1; ‘landmark tree’) in spring 2012. The samples comprised eight twigs from the crown at four heights (3, 8, 13 and 18 m) and two orientations (north and south), and two trunk cores (same orientations). Cores were extracted using a sterilised core drill. The 25-m tree was a lingering monumental elm. Common garden tests on clones generated from its cuttings showed that the tree was not genotypically resistant to DED (data not shown), and in 2014 it died after an exceptionally harmful DED outbreak.

**Fig. 1.**
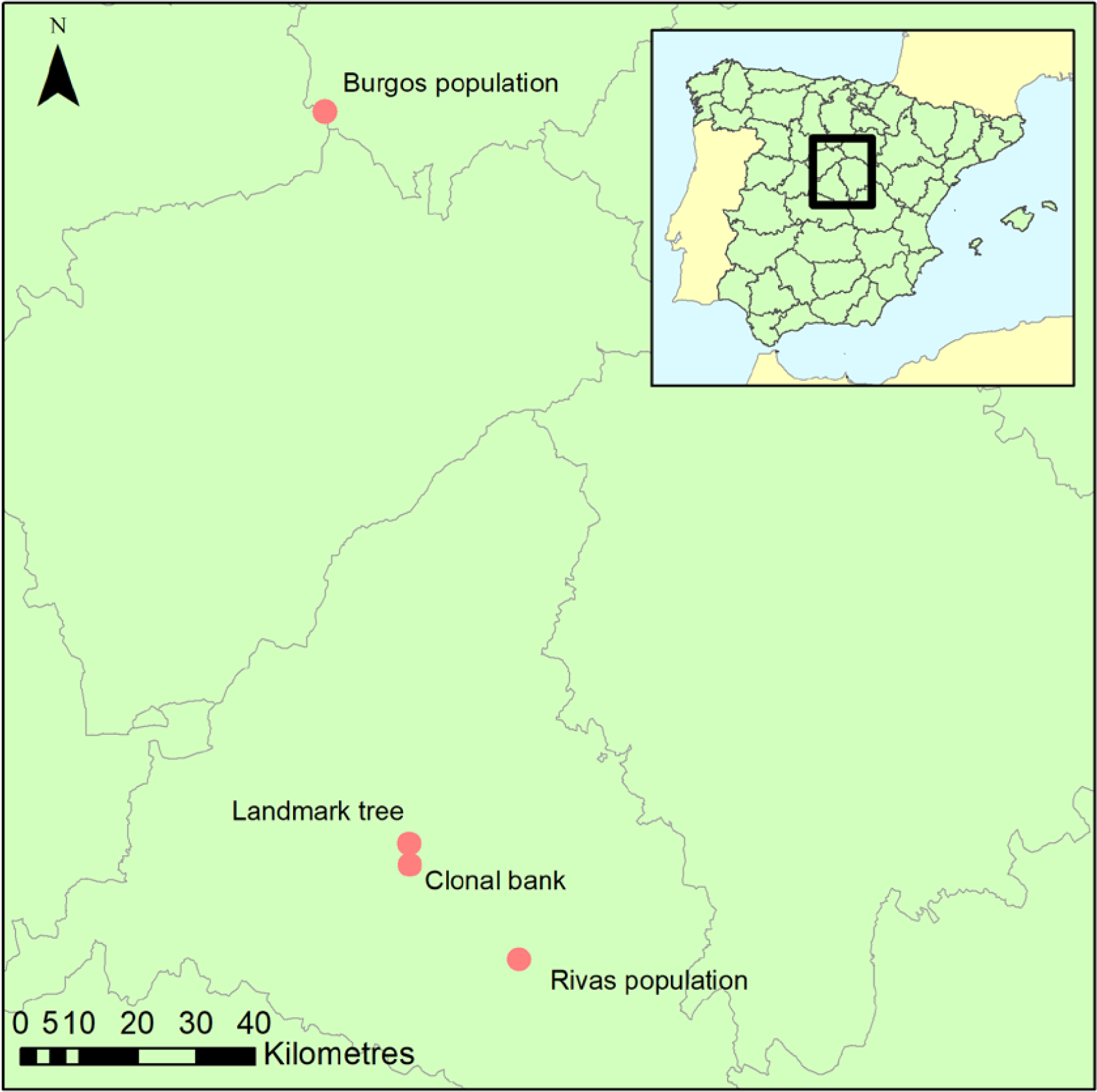
Geographical location of the four sample collection sites.

#### ii) Wood mycobiome and elm DED tolerance

The second sampling was at the elm clonal bank (common garden setup) at the *Puerta de Hierro* Forest Breeding Centre (Madrid; Fig. 1), the headquarters of the Spanish elm breeding programme. The clonal bank has around 250 genotypes from Spain, including seven DED-tolerant genotypes [19]. Four twigs were collected from scaffold branches in 10 trees (Sup. Table 1) catalogued as tolerant (n=3; V-AD2; M-RT1.5; M-DV5), intermediately susceptible (n=4; CR-RD2; GR-HL2; J-CA2; MA-PD2), or susceptible (n=3; GR-DF3; M-DV1; TO-PB1). Samples were collected at four locations per tree to ensure accurate representation of the endophyte composition and mitigate any effect of local infections (see below). All twigs were collected from the lower half of the crown, to a height of 4 m.

The level of tolerance to DED of the 10 *U. minor* clones sampled at the clonal bank was determined during screening tests at the Spanish elm breeding programme at *Puerta de Hierro* Forest Breeding Centre (Madrid, Spain) (Sup. Table 1, Sup. Text). The 10 trees sampled were not inoculated with the pathogen.

#### iii) Variation in trees differing in vitality phenotype

Following the same protocol as at the clonal bank, twigs from six trees were collected from a natural *U. minor* stand in the municipality of Rivas-Vaciamadrid (Rivas population; ‘Madrid province’; Fig. 1). This population lacks genotypically tolerant clones (tested in a common garden) but has not been destroyed by DED. The reasons for its survival are unclear but could be explained by phenotypical tolerance due to the effect of biotic or abiotic factors. The stand nonetheless shows clear signs of dieback, due in part to DED infections but also to undetermined causes. We collected samples from trees representing phenotypes ranging from healthy to various stages of dieback (Sup. Fig. 1). The health status of the sampled trees (named RV1 to RV6) was estimated visually, using a score from 1 (no symptoms) to 6 (profuse dieback symptoms).

#### iv) Variation among geographical locations

Using the same protocol as at the clonal bank, three trees from a small, young natural stand in the province of Burgos (approximately 150 km north of the other locations; Fig. 1) were sampled to provide a background reference of endophyte diversity for the populations in Madrid province using a distant population.

### DNA isolation, amplification and NGS

After collection, samples were sterilised, frozen for storing, and ground. The individual tree samples from the clonal bank, Rivas and Burgos populations were pooled and milled together, resulting in only one tube of wood powder from each tree containing a mixture of the four twigs collected. DNA was isolated from the powder after enzymatic digestion to improve recovery of fungal DNA. Zirconium oxide beads were added during vortexing to increase cell wall lysis. Endophyte composition was profiled by high throughput sequencing of the first internal transcribed spacer region (ITS1) of the ribosomal DNA. Sequencing effort was uneven among experiments, prioritising the landmark tree samples, which was also the first processed, to determine the level of resolution needed in subsequent experiments. The clonal bank experiment followed in sequencing effort, to attain accurate values of endophyte abundance for identifying associations with DED tolerance. The Burgos population was only shallowly sequenced due to its outgroup condition, to reveal ubiquity of microbiome elements detected in the other populations. DNA amplification was performed in two steps: (1) to cover the target region with oligonucleotides that contained the specific fungal primer ITS1-F [20] or the non-specific primer ITS2 [21]; (2) to attach the adaptors for the sequencing platform. After the second PCR, the product of all the samples was quantified, pooled equimolarly and pyrosequenced in a 454 GS FLX Titanium platform (Roche, Basel, Switzerland).

### Bioinformatic pipeline

The bioinformatic treatment of pyrosequencing output was performed following the guidelines of Lindahl et al. [22]. Demultiplexing, denoising, dereplication, dechimerisation and sequence truncation processes were carried out using the default values of the *RunTitanium* script developed in AmpliconNoise v1.29 [23] The ITS1 region was then extracted from the sequences using FungalITSextractor [24].

Although AmpliconNoise creates OTUs (Operational Taxonomic Units) by collapsing identical sequences, we further clustered them with the grammar-based software GramCluster 1.3 [25] in greedy mode to build new OTUs, allowing higher variation among sequences. This programme was run on the whole dataset (i.e. pooling the output of all samples) to build OTUs across all samples, allowing subsequent among-sample comparisons.

### Taxonomic assignment

Because taxonomic composition was of paramount interest, a complex methodology was used to assign taxonomy to OTUs. Sequence similarity to a curated database [26, 27] (UNITE) was the first approach. Because a large number of OTUs had undetermined taxonomies, an in-house method was used to further assign the higher taxonomic ranks, restricting haplotype identity to the very well conserved regions of the ITS1 and small flanking regions.

### Diversity estimates and hypothesis contrasts

Commonly used diversity indices were estimated for each sample collected, using the counts per OTU as taxonomic information. Simpson’s and Shannon’s indices, and species richness on counts rarefacted to 500, were calculated using R package “vegan” v. 2.3-0 [28]. Statistical analyses were performed taking into account that count data in these types of studies follow a negative binomial distribution as in RNA-seq experiments [29]. As suggested by these authors, R package DESeq2 [30] was used to examine the data and test for associations between taxonomic group abundance and tolerance to DED. Given the unreliable taxonomic certainty of OTU formation through clustering and the possible redundancy in ecological function of closely related species and genera, we preferred to focus on the higher taxonomic levels.

### Core microbiome demarcation

The distributions of number of samples in which each OTU was present (OTU incidence distribution) were used to determine which OTUs were putatively from the core microbiome. The expected pattern of incidence of OTUs, if their occurrence probability is low and mostly based on randomness (i.e. local infections rather than core microbiome), must agree with a Poisson or negative binomial distribution. OTUs present in at least eight of the 10 samples from the landmark tree or the clonal bank were considered to be members of the core microbiome. We selected eight samples as the threshold value because this is where the distributions clearly diverge from Poisson distributions (see Results).

## Results

### Sampling effort and saturation

After running the bioinformatic pipeline, we obtained 106,047 informative reads (considered counts). These were grouped by GramCluster into 435 clusters (considered OTUs henceforth). Of these, 74 were singletons, 40 doubletons and 23 tripletons. A further 263 OTUs were represented by more than five reads. To ensure a more accurate OTU richness comparison, we rarefied the count data to 500 reads per sample. The mean values (± s.e.) of rarefied OTUs ranged from 66.0 ± 4.0 in one of the lower resprouted branches of the Somontes tree to 16.7 ± 2.3 in one sample from the Rivas stand (RIV2, with advanced dieback). Rarefaction curves supported the figures observed by the rarefaction to 500 reads and indicated that the sampling effort was sufficient to capture the richness trends of each sample (Sup. Fig. 2 and 3). Principal Component Analysis showed a separation between sites (Fig. 2).

**Fig. 2.**
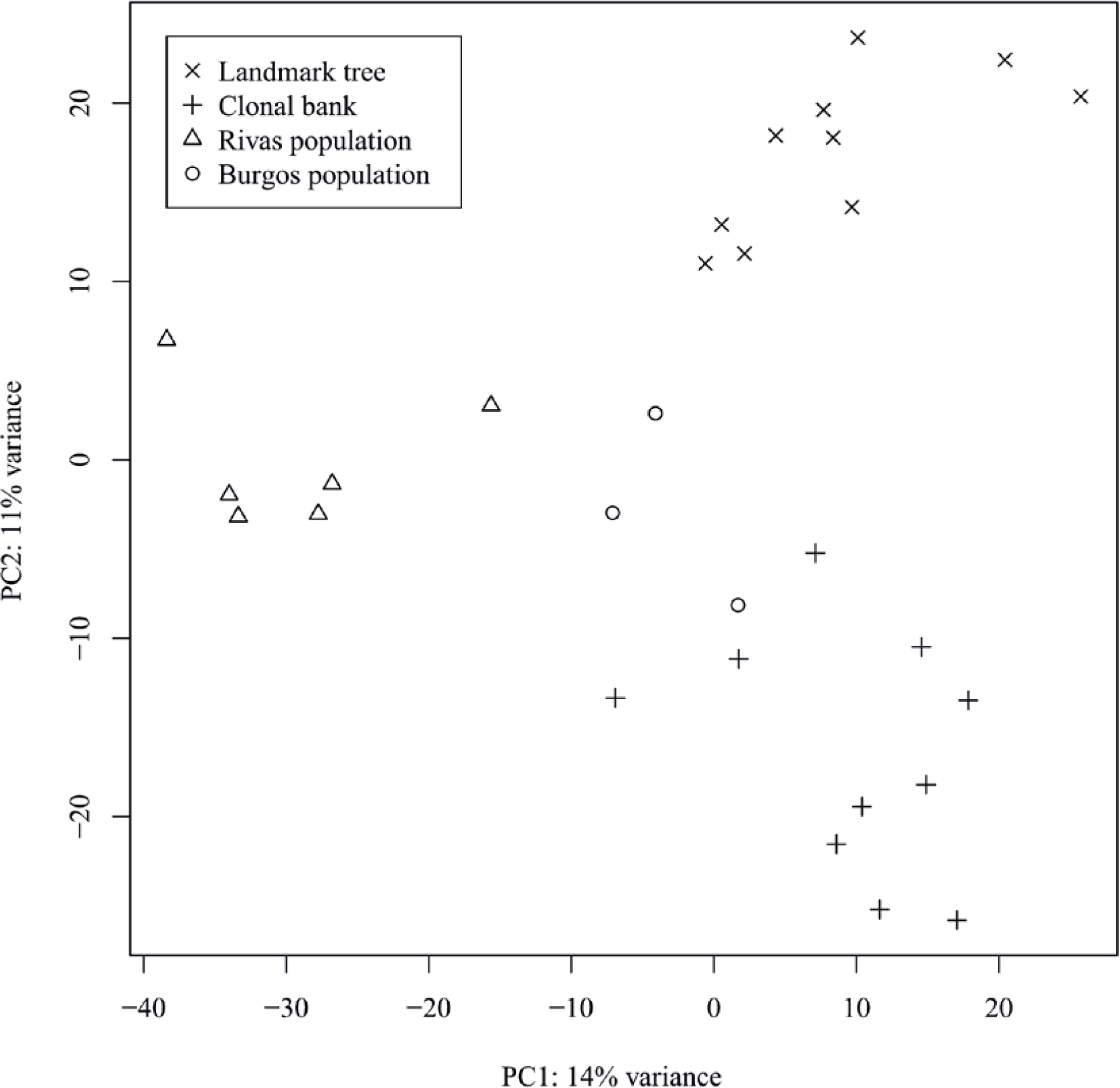
First two axes from the Principal Component Analyses performed on the OTU counts after variance stabilising transformation.

Across the total sample set, 96 families, 47 orders, 19 classes and 4 phyla were detected. Of the 435 OTUs detected, 427 were assigned to a phylum, 352 to a class, 301 to an order and 219 to a family. Genus was provided for 206 OTUs, and species for 145.

Applying the procedure of screening highly conserved runs of the ITS1 and flanking sequences, we were able to infer the identity of 75 of the 83 OTUs whose phyla were unidentified using only a BLAST search on the UNITE database (further information in Sup. Results).

### Within-tree distribution of endophytes

The Somontes tree had 69,558 reads passing filtering, clustered into 284 OTUs (39 singletons, 20 doubletons and 14 tripletons). Regarding incidence, 11 OTUs were present in all the in-tree locations sampled and 23 were present in at least eight (Table 1; Fig. 3a). A further 129 OTUs were present in just one location and 60 were present in two (Fig. 3a). The number of OTUs at higher abundance in the tree did not follow a purely rare event distribution such as the Poisson or negative binomial distribution, as seen in the smooth but distinguishable peak at the end of the distribution (Fig. 3a). Four phyla, 17 classes, 38 orders and 76 families were detected within the tree (Fig. 4). Across the tree, the levels of diversity (measured as Shannon’s H, Simpson’s λ and rarefied OTU richness) were generally high, with the following deviations: (i) the two lowest branches, produced from resprouts from the trunk, displayed remarkably higher levels of diversity; (ii) one sample from the trunk and one from the middle crown exhibited low values of both H and λ.

**Table 1.**
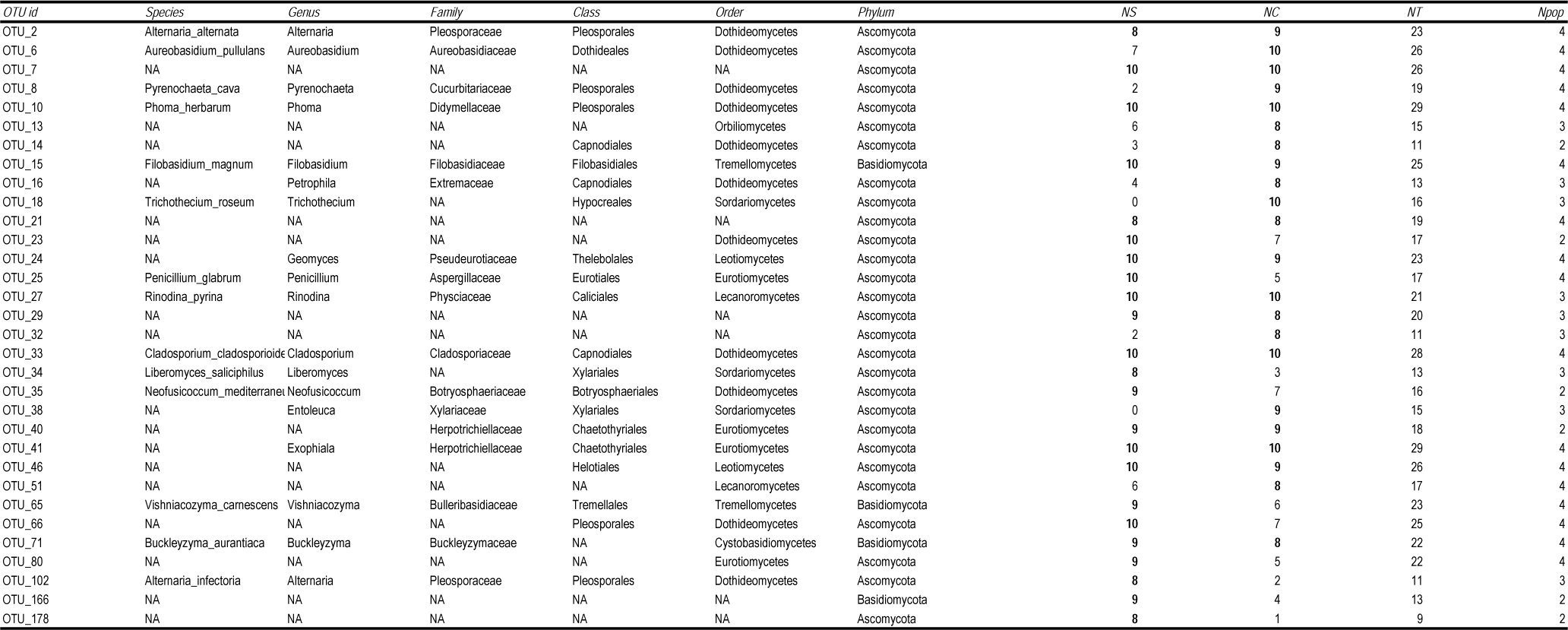
OTUs present in at least eight samples of the landmark tree or the clonal bank. Taxonomic assignment is based on ITS1 DNA similarity with UNITE database, with some in-house modifications. The final columns show the number of samples in the landmark tree (*NS*), the clonal bank (*NC*) and the total sample set (*NT*) and the number of geographical locations (*Npop*) where the OTUs were detected.

**Fig. 3.**
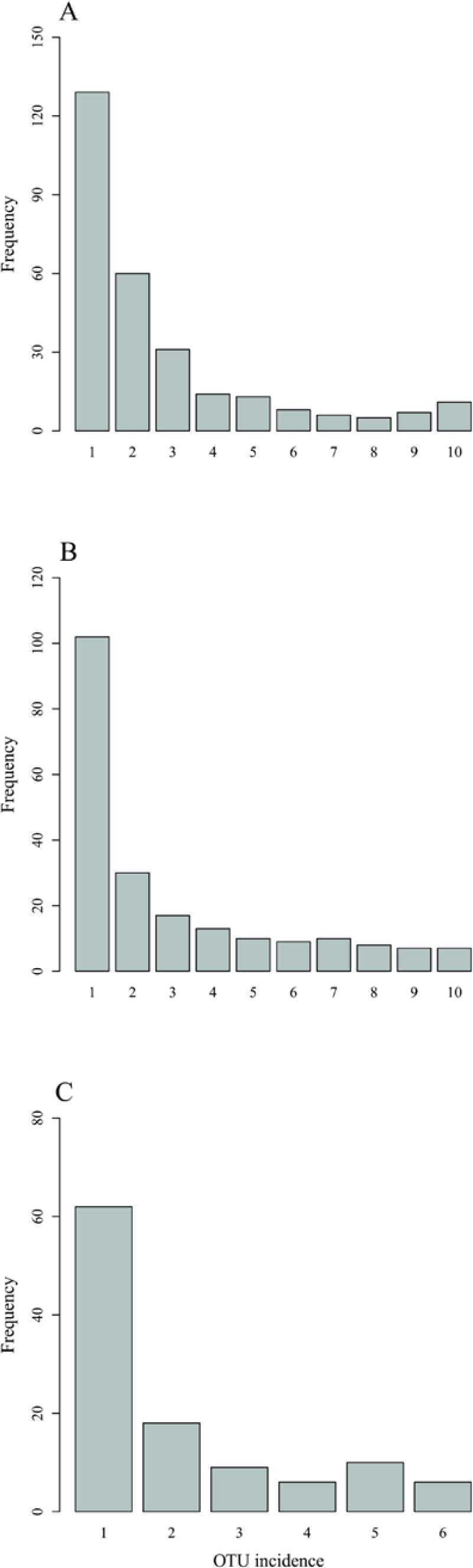
OTU frequency spectra for (A) landmark tree, (B) clonal bank and (C) Rivas population.

**Fig. 4.**
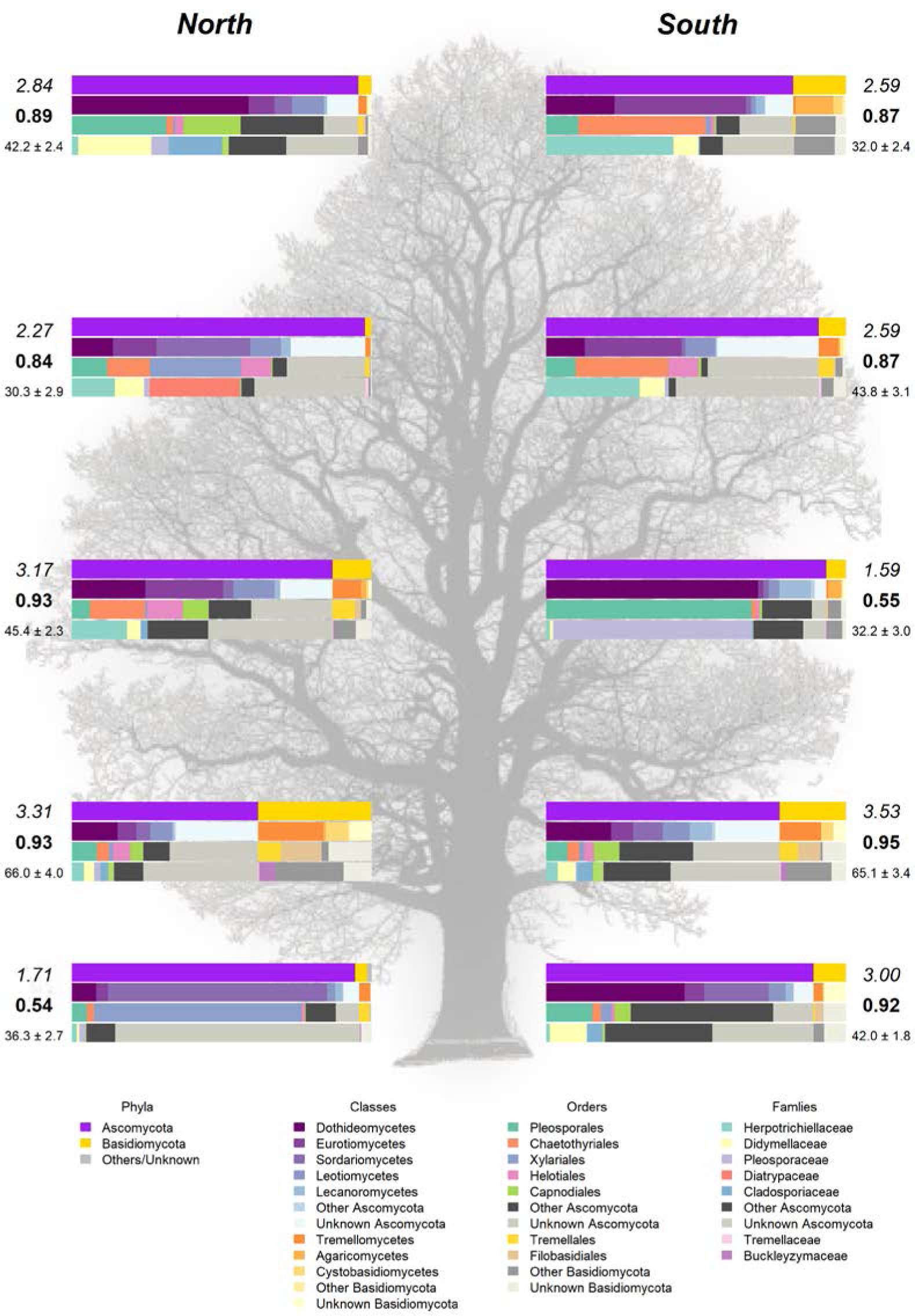
Taxonomic composition in the landmark tree. Only the most relevant taxa are shown. Coloured bars represent the frequency of taxa at the levels of phylum, class, order and family (top to bottom). Numbers next to the bars indicate the Shannon (italics) and Simpson (bold) indices and the OTU richness rarefacted to 500 reads (with standard error). (Background image source: Tree Silhouette copy by Bob G in flickr, licensed under CC BY-NC-SA 2.0).

### Endophyte diversity in relation to DED tolerance

High-throughput sequencing on the 10 trees of varying levels of tolerance to DED from the clonal bank at *Puerta de Hierro* research centre produced 20,613 sequences after filtering. The sequences were clustered into 213 OTUs: 49 singletons, 20 doubletons and 19 tripletons. Similar to the results in the Somontes tree, most OTUs were present in just one sample (102), two samples (30) or three samples (17). However, the counts did not drop at a rate consistent with a Poisson process, and reached a stable level beyond five samples (Fig. 3b). In total, two phyla, 15 classes, 29 orders and 61 families were detected (Fig. 5a).

**Fig. 5.**
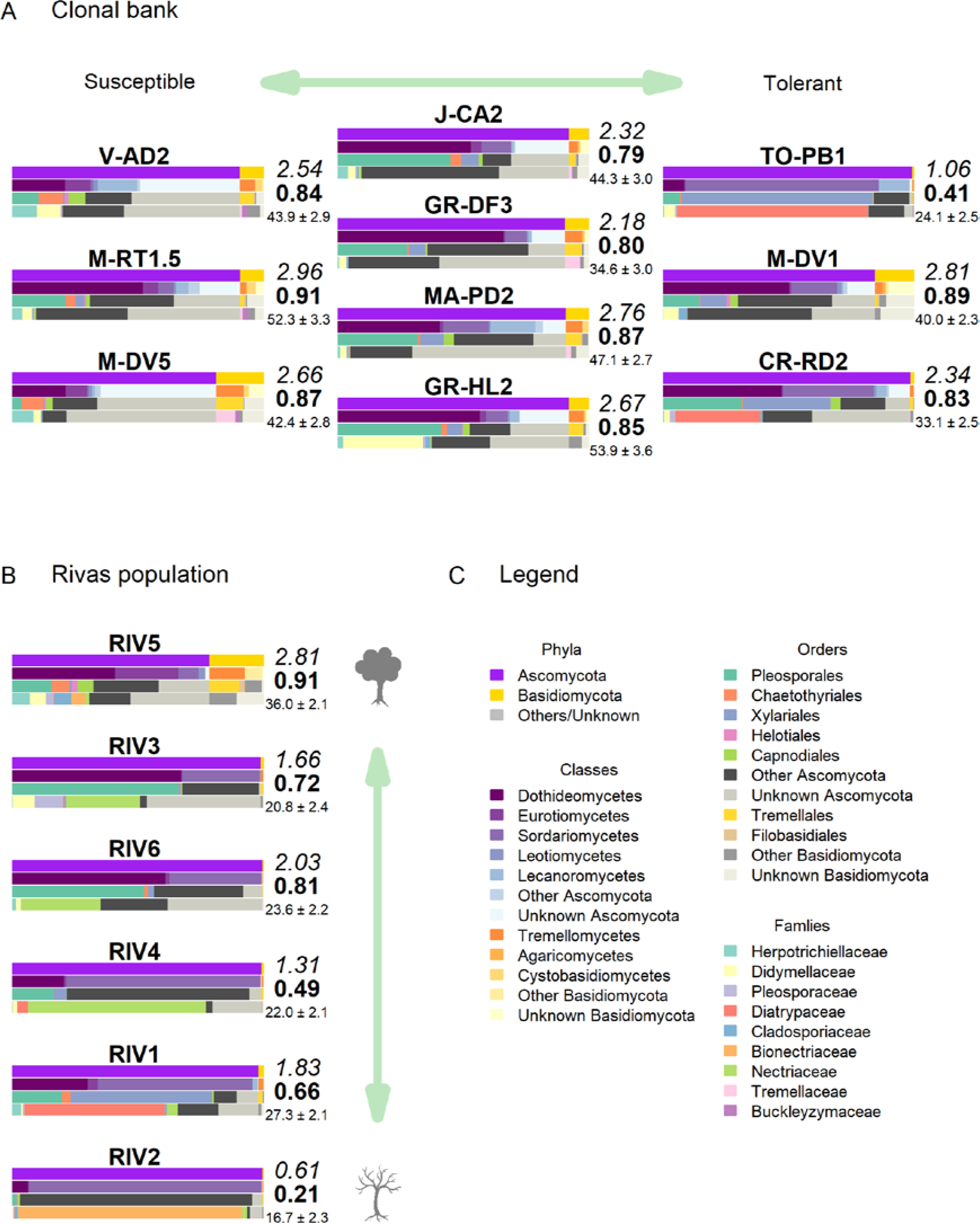
Taxonomic composition in (A) clonal bank and (B) Rivas population. Only the most relevant taxa are shown. Coloured bars represent the frequency of taxa at the levels of phylum, class, order and family (top to bottom), following legend color code (C). Numbers next to the bars indicate the Shannon (italics) and Simpson (bold) indices and the OTU richness rarefacted to 500 reads (with standard error). (Tree icon sources: minimal tree simple SVG Silh, licensed under CC0 1.0 and tree-304418 by Clker-Free-Vector-Images in pixabay under Pixabay licence).

Clone TO-PB1 (susceptible) displayed the lowest levels of diversity. Conversely, the resistant clone M-RT1.5 showed the highest overall diversity estimates. GR-HL2 (susceptible) and MA-PD2 (moderately tolerant) also displayed high diversity values. Wilting after DED inoculation (used as a proxy of susceptibility) was not significantly correlated with any of the diversity estimates, indicating the absence of a strong correlation between diversity estimates and tolerance to DED. However, the limited sample size (*n* = 10) may have prevented detection of a more subtle correlation.

The tests of association between wilting and taxa abundance produced unambiguous hits (Table 2). Three families and three orders were significantly associated with tolerance. The family with the highest association was Buckleyzymaceae (Fig. 6a), a Basidiomycota of the Cystobasidiomycetes class and undefined order. Because the order was undefined, this association was reproduced at class level rather than order level. It had lower support at OTU level, represented by the genus *Buckleyzyma* (OTU_71). The next most significant hit was from the family Herpotrichiellaceae, Ascomycota (Fig. 6b). This is the only family of the order Chaetothyriales and class Eurotiomycetes detected in the study and therefore the hit was reproduced at these taxonomic levels. It was also supported, but to a lesser degree, by the hit at OTU level, in OTU_41 assigned to the genus *Exophiala*. The next and least significant hit at family level was Tremellaceae (Fig. 6c), echoing at order level as Tremellales (Basidiomycota). Two OTUs were significant to a lesser degree and belonged to the genus *Cryptococcus* (Table 2). All these taxa were negatively associated with susceptibility (proxied as wilting). At order level, the Xylariales were positively associated with susceptibility, and this result was reproduced with similar support at class level (Sordariomycetes), partly because the Xylariales are the most abundant group in Sordariomycetes. The positive association of OTU_19 (Sordariomycetes), without further taxonomic assignment, hints at a general relationship between the Sordariomycetes and susceptibility.

**Table 2.** Taxa with significant positive or negative associations (padj < 0.1; *p* -value < 0.05 for OTUs) with tolerance to DED. The test of association was performed by a Wald test. Column *baseMean* shows the mean of normalised counts; *log2FC* : estimate of the effect size scaled to the log2 of fold change; *lfcSE* : standard error of this estimate; *stat* : value of the Wald test statistic; and *p-value* and *padj* : respectively, the raw and the adjusted (for multiple tests) probabilities that the observed statistic is part of the null distribution. These columns correspond to the output of the function DESeq from R package DESeq2. A positive fold change indicates association with susceptibility to DED.

| Taxon | *baseMean* | *log2FC* | *lfcSE* | *stat* | *p-value* | *padj* | *Genus* | *Class* |
| --- | --- | --- | --- | --- | --- | --- | --- | --- |
| Class |  |  |  |  |  |  |  |  |
| Sordariomycetes | 796.4585 | 0.075371 | 0.016651 | 4.526652 | 5.99E-06 | 8.39E-05 |  |  |
| Cystobasidiomycetes | 18.31421 | -0.05685 | 0.014337 | -3.96537 | 7.33E-05 | 0.000513 |  |  |
| Eurotiomycetes | 57.25089 | -0.02946 | 0.010795 | -2.72913 | 0.00635 | 0.029634 |  |  |
| Order |  |  |  |  |  |  |  |  |
| Xylariales | 567.7225 | 0.076165 | 0.017111 | 4.451289 | 8.54E-06 | 0.000213 |  |  |
| Chaetothyriales | 76.75799 | -0.04752 | 0.014288 | -3.32587 | 0.000881 | 0.011018 |  |  |
| Tremellales | 93.79415 | -0.03451 | 0.012984 | -2.65755 | 0.007871 | 0.065592 |  |  |
| Family |  |  |  |  |  |  |  |  |
| Buckleyzymaceae | 14.7488 | -0.07458 | 0.018857 | -3.95538 | 7.64E-05 | 0.003362 |  |  |
| Herpotrichiellaceae | 56.56981 | -0.04241 | 0.012086 | -3.50896 | 0.00045 | 0.009897 |  |  |
| Tremellaceae | 48.44816 | -0.06776 | 0.023945 | -2.82967 | 0.00466 | 0.06834 |  |  |
| OTU |  |  |  |  |  |  |  |  |
| OTU_70 | 19.43169 | -0.11353 | 0.036497 | -3.11056 | 0.001867 | 0.141228 | Cryptococcus | Tremellomycetes |
| OTU_71 | 13.33833 | -0.07047 | 0.024342 | -2.8951 | 0.00379 | 0.141228 | Buckleyzyma | Cystobasidiomycetes |
| OTU_55 | 20.02071 | -0.11923 | 0.041255 | -2.89006 | 0.003852 | 0.141228 | Cryptococcus | Tremellomycetes |
| OTU_19 | 4.591395 | 0.078549 | 0.028191 | 2.786321 | 0.005331 | 0.146603 | NA | Sordariomycetes |
| OTU_41 | 49.44945 | -0.04613 | 0.01862 | -2.47718 | 0.013243 | 0.291339 | Exophiala | Eurotiomycetes |

**Fig. 6.**
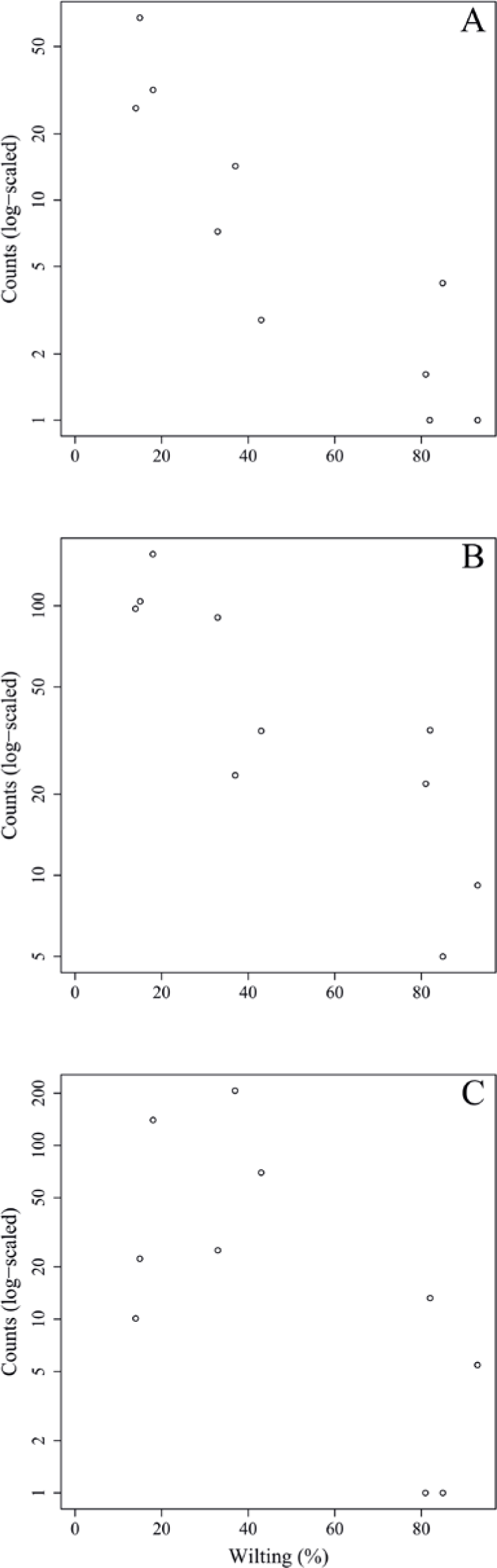
Relation between susceptibility to DED (measured as leaf wilting percentage) of the 10 clonal bank genotypes and the normalised counts detected from reads of endophytic fungal families (A) Buckleyzymaceae, (B) Herpotrichiellaceae and (C) Tremellaceae.

### Endophytic mycobiome in trees representing a gradient of vitality

The six samples collected in the natural riparian stand at Rivas-Vaciamadrid municipality from trees at varying stages of dieback produced 13,432 reads, clustered into 111 OTUs: 30 singletons and 16 doubletons. Forty-eight were represented by more than five reads. Only six OTUs were present in all trees and 10 were present in five samples (Fig. 3c). The secondary peak found in the OTU incidence distribution was not in the total number of samples (*n* = 6) but in *n* = 5.

None of these OTUs was identified as genus *Ophiostoma* or order Ophiostomatales, even though the UNITE database included several accessions for both *O. ulmi* and *O. novo-ulmi*. The most affected tree (RIV2) and two trees with moderate dieback (RIV1 and RIV4) were dominated by Sordariomycetes: RIV1 was rich in Diatrypaceae and RIV2 in Bionectriaceae (Fig. 5b). Both RIV4 (moderate dieback) and RIV6 (incipient dieback) had Nectriaceae as the most abundant family, although it was also abundant in the healthy RIV3. The two healthy trees (RIV3 and RIV5) were more infected than the other trees by Dothideomycetes and Eurotiomycetes. For diversity, RIV5 exhibited the highest values in all three indices calculated (Shannon’s H, Simpson’s λ and rarefied OTU richness). The affected RIV1 and RIV6 displayed high values of H and richness, and RIV3 (healthy) and RIV6 had high values of λ. The tree with lowest vitality (RIV2) had the lowest diversity values.

The healthiest tree (RIV5) displayed a clearly distinct pattern that was much richer in Basidiomycota (Fig. 5b). Aureobasidiaceae was the most common family in this tree, followed by Herpotrichiellaceae. The microbiome of RIV2, a tree with low vitality (RIV2), was dominated by Bionectriaceae (OTU_147, genus *Geosmithia*). This OTU was virtually absent in the other samples, except in the healthiest (RIV5), where it was not abundant but had a significant presence.

Regarding the taxa significantly associated with DED tolerance, Buckleyzymaceae (represented mostly by OTU_71) was virtually absent from the population. Herpotrichiellaceae (represented mostly by OTU_41) was present in all trees but was much more abundant in RIV1 (dieback) and RIV5 (very healthy). Tremellaceae was present in RIV4 and marginally in RIV6. The single OTU associated with increased DED susceptibility (OTU_19; Sordariomycetes) was detected only marginally in RIV1 (dieback).

### Patterns across the four sites – core fungal endobiome of U. minor

To determine the extent of ubiquity of the most common OTUs, we examined the incidence of OTUs in the global sample set (*n* = 29) by pooling all samples analysed. Of the 435 OTUs detected, 192 were present in only one sample, 76 in two samples and 34 in three samples. Distribution then reached a local maximum at six samples (Sup. Fig. 4). Two clusters were present in all 29 samples (OTU_10, Didymellaceae, Dothideomycetes; and OTU_41, Herpotrichiellaceae, Eurotiomycetes, associated with DED tolerance, see above), one was present in all but one (OTU_33, Cladosporiaceae, Dothideomycetes), and three others were present in all but two (Table 1). Beyond the category of “presence in nine samples” distribution was effectively flat. In other words, the number of OTUs present in 10 to 29 samples always ranged from 1 to 5 (Sup. Fig. 4). A further 37 OTUs were found in the four sampled populations, 44 in three, 90 in two, and 264 were private to a single population. These two distributions concur with the distributions of OTUs in the clonal bank and, to a lesser extent, with that of the OTUs in the Somontes tree. The OTUs present more frequently in our sampling than could be expected by chance are very likely members of the core microbiome (see Discussion). In total, 32 OTUs passed the criteria for core microbiome membership: 28 belonging to Ascomycota and four to Basidiomycota.

## Discussion

### Within-tree variation in species richness and diversity

Analysis of the landmark tree endophytic mycobiome revealed no conspicuous pattern, but it was possible to draw some conclusions: (i) although most of the samples collected displayed a similar taxonomic composition, some were remarkably different, e.g. one from the southern mid-height branch (H1S), which was massively infected by a single OTU (Fig. 4); (ii) the two lowest branches, resprouts from the trunk aged only a few years (epicormic shoots), displayed higher taxonomic richness than the other branches, with a relatively high representation of Basidiomycota; and (iii) samples from the main trunk showed a richness comparable with that of the crown branches. Taking this into consideration, when sampling trees to characterise their overall stem endophytic flora and to avoid considerable bias due to abnormally high local infection, pooling tissue from at least two branches is recommended. However, mixing samples from epicormic and crown branches should be avoided, because they are likely to represent different endobiome compositions. The greater richness found in the lower branches supports previous research [6, 31] and could be partly attributed to the high density of airborne inoculum near the ground. Similarly, as a substrate for fungi, epicormic shoots may differ in anatomy and quality from proleptic shoots [32].

### Endobiome and tolerance to DED

The abundance of three distinct fungal endophytic taxa was associated with higher host tree tolerance to DED. Interestingly, the two highest associations at family level (Buckleyzymaceae, in Cystobasidiomycetes; Herpotrichiellaceae, in Eurotiomycetes) were mostly driven by OTUs considered to be members of the core microbiome (OTU_71 and OTU_41, respectively). Moreover, a common trait of these three taxa associated with DED tolerance is that they grow, or are able to grow, as yeasts. Yeasts have the ability to systemically colonise plants and produce phytohormones and siderophores that promote plant growth and alleviate stress [33]. The greater abundance of these yeasts in low-susceptible trees could improve tree resilience to DED infection, promoting tree tolerance mechanisms to the physiological disorders caused by the pathogen. *Ophiostoma novo-ulmi* also spreads systemically through the plant’s vascular system in a yeast-like phase [34] (blastospores), even in tolerant trees [35], inducing vessel cavitation. Our results suggest that tolerance to DED is enhanced in trees harbouring a high abundance of two yeasts from the core endobiome (OTU_71 and OTU_41), which have the capacity to extensively colonise the plant. Extensive or systemic spread of an endophyte could allow higher interaction with the pathogen throughout the plant, and possibly a higher level of interaction with the plant’s physiological functions.

The first endophyte was assigned to *Buckleyzyma aurantiaca*, based on the sequence similarity to the accessions in the database UNITE. When the ITS sequence of this OTU was run against Genbank, equal hits were returned for several accessions identified as *Buckleyzyma* and *Rhodotorula*, both cultured and uncultured, but with a level of identity of 97.22% (140/144 bp). This OTU is likely to be an undescribed species. Cystobasidiomycetes is a group of basidiomycetous yeasts with unclear systematics that includes strains previously isolated from plants [36], soils and waters [37–39].

The second endophyte (OTU_41) was assigned to *Exophiala* by our pipeline. In Genbank, it did not retrieve perfect identities, obtaining a maximum identity of 97.55% (196/201 bp) and three gaps. Most accessions were derived from uncultured strains, and some from molecular studies in soils and plants. This OTU could therefore also belong to an undescribed species. The Herpotrichiellaceae belong to the black yeasts and have been reported to grow in the sexual phase in dead plants and wood [40].

The third associated taxon was represented by two OTUs (OTU_70 and OTU_55) of the genus *Cryptococcus* (Tremellomycetes), another yeast frequently found in plants and water [39]. Albrectsen et al. [41] found *Cryptococcus* as an endophyte in beetle-damaged *Populus tremula* leaves, and *Rhodotorula* in the beetles. In addition, *Cryptococcus* apparently outcompetes the Rosaceae pathogen *Botrytis cinerea* due to niche occupancy [42].

### Phenotypic vitality and wood mycobiome

Some conspicuous taxa showed up in the tree vitality analysis. Similar to our results (in the declining tree RIV2), *Geosmithia* spp. was identified as the dominant fungi in a *U. minor* tree with extensive dieback symptoms in the absence of DED pathogens [43]. Certain *Geosmithia* fungi could therefore act as opportunistic or latent pathogens in elms. The presence of this genus in the healthiest tree (RIV5) suggests that it is able to live as an endophyte in latent pathogenicity. Pepori et al. [44] found that elms inoculated with *Geosmithia* fungi remained largely asymptomatic, and joint inoculation of *Geosmithia* and *O. novo-ulmi* reduced wilting symptoms compared to inoculation with *O. novo-ulmi* only. They also found parasitic behaviour of *Geosmithia* towards *O. novo-ulmi*. In elms, *Geosmithia* was frequently found in DED-infected trees [45], most likely carried there by the beetles that are also the vectors of DED pathogens. Further research is needed into the potential contribution of *Geosmithia* to tree dieback in Rivas or, in contrast, the potential role of this taxon in the phenotypic tolerance to DED found in this elm stand.

Two other trees with dieback symptoms (RIV6 and RIV4) were dominated by Nectriaceae (especially RIV4). OTU_92 (unknown genus) was responsible for this signature and was also very abundant in the healthy RIV3. The family Nectriaceae includes facultative parasites that cause stem cankers, and saprobes. In elms, dieback symptoms have been associated with colonisation by *Nectria* sp. [46, 47].

### Core microbiome and among-site variation

Sampling from different locations in a single tree and from genotypically different trees enabled detection of robust signatures of a core microbiome, following the concept of Shade and Handelsman [3]. Of the 245 nonsingleton OTUs found in the landmark tree, 11 were present in all samples (10) and 23 in more than seven samples (Table 1). In the clonal bank, eight OTUs were present in eight trees, seven were present in nine trees and another seven were in all trees (10). In the landmark tree and the clonal bank, the number of OTUs did not decrease following the pattern expected by randomness. The number of OTUs reached a tableau beyond five samples in both distributions (Fig. 3a, b), and a relative maximum at the end of the distribution in the landmark tree (Fig. 3a). Therefore, the probability that a given sample would contain a specific OTU depended on the OTU in question. Not all OTUs can be considered rare events (i.e. events that would retrieve Poisson distributions). Others with high probabilities of occurrence displayed different distributions (Poisson distributions, but with “absence of OTU” as rare event). Although not appreciable, perhaps due to their low numbers, other OTUs may have behaved as “medium frequency events”, retrieving binomial distributions. Thus the lack of agreement between the observed distributions and the expected monotonic decrease, characteristic of pure Poisson processes, shows that OUT occurrences range from rare to highly frequent. OTUs that follow a pattern of occurrence consistent with a Poisson distribution could be considered local infections with arguably different but low likelihoods of infecting a stem. Highly frequent OTUs, on the other hand, are likely to be members of the core microbiome. It is unclear why this latter group of endophytes is pervasive, but it could be explained by a high infective capacity [48] (e.g. through insect vectors, rain and wind) and/or systemic propagation within the plant, as occurs in some endophytic yeasts [33]. Shallower sampling may not have allowed us to distinguish between the two trends in OTU occurrence, because the distributions would have overlapped, obscuring the underlying pattern. The most commonly found fungal taxa both in the landmark tree and the clonal bank were the ascomycetous classes Dothideomycetes, Eurotiomycetes, Sordariomycetes, Leotiomycetes and Lecanoromycetes, and the basidiomycetous classes Tremellomycetes and Cystobasidiomycetes.

Defining the core microbiome as the OTUs that are present in at least eight out of 10 samples in either the landmark tree or the clonal bank, we identified 32 core OTUs. Although most of them were present in most samples across the four populations, some were abundant in the clonal bank but rare or absent in the landmark tree (e.g. OTU_18 and OTU_38). Considering that the clonal bank includes trees from various provenances across Spain (Sup. Table 1) and a few are from the same provenance as the landmark tree, it is conceivable that these OTUs are controlled mostly by environmental cues [9]. Conversely, a few OTUs were widespread in the landmark tree, but rarer in the clonal bank (e.g. OTU_66, OTU_80 and OTU_102). OTU_66 and OTU_80 were present in the four populations and most of the samples but surprisingly lacking in some trees from the clonal bank. This pattern hints at an implication of host genotype (see Bálint et al. [49]). However, physiological status and microscale environmental variation could also explain this pattern. The clear separation of samples by site shown in the Principal Component Analysis (Fig. 2) indicates the important role of geographical location in shaping fungal endobiome communities. New targeted experiments are needed to confirm or refute these hypotheses.

## Acknowledgements

This work would have not been possible without the support of the staff of the Spanish elm breeding programme (Spanish Ministry of Agriculture, Fisheries and Food): Jorge Domínguez Palacios, David León Carbonero, Felipe Pérez Martín and Salustiano Iglesias. Funding for this work was provided by the Spanish National Plan for Scientific and Technical Research and Innovation, Spanish Ministry of Economy and Competitiveness, through project AGL2015-66925-R (MINECO/FEDER).

## Competing Interests

The authors declare no competing financial interest in relation to the work described.

## Notes

### Competing Interest Statement

The authors have declared no competing interest.

