## Supplementary material for "Structure of core fungal endobiome in *Ulmus minor*: patterns within the tree and across genotypes differing in tolerance to Dutch elm disease"

**Supplementary Information**

Sup. Fig. 1. Pictures of trees from the Rivas natural population.

Sup. Fig. 2. Rarefaction curves for the 29 samples collected.

Sup. Fig. 3. Rarefaction curves grouped by collection site.

Sup. Fig. 4. OTU frequency spectra for total sample set.

Sup. Table 1. Susceptibility to DED in the trees of the clonal bank.

Sup. Text 1. Detailed methods.

Sup. Text 2. Supplementary results on sequencing effort, taxonomic assignment and taxonomical composition.

Sup. Text 3. Taxonomical considerations.

Sup. Text 4. Supplementary references.

A) RIV1

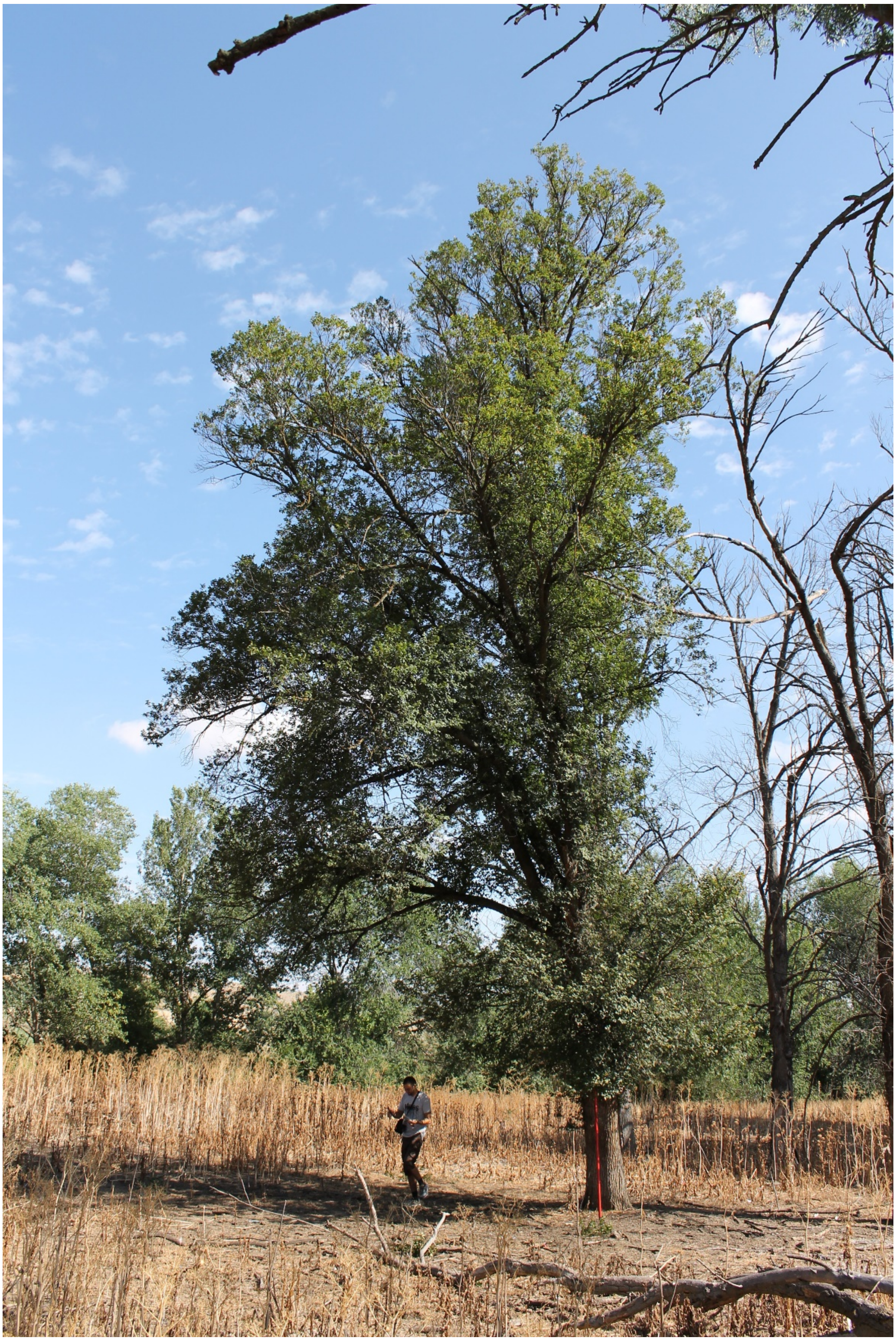

B) RIV2

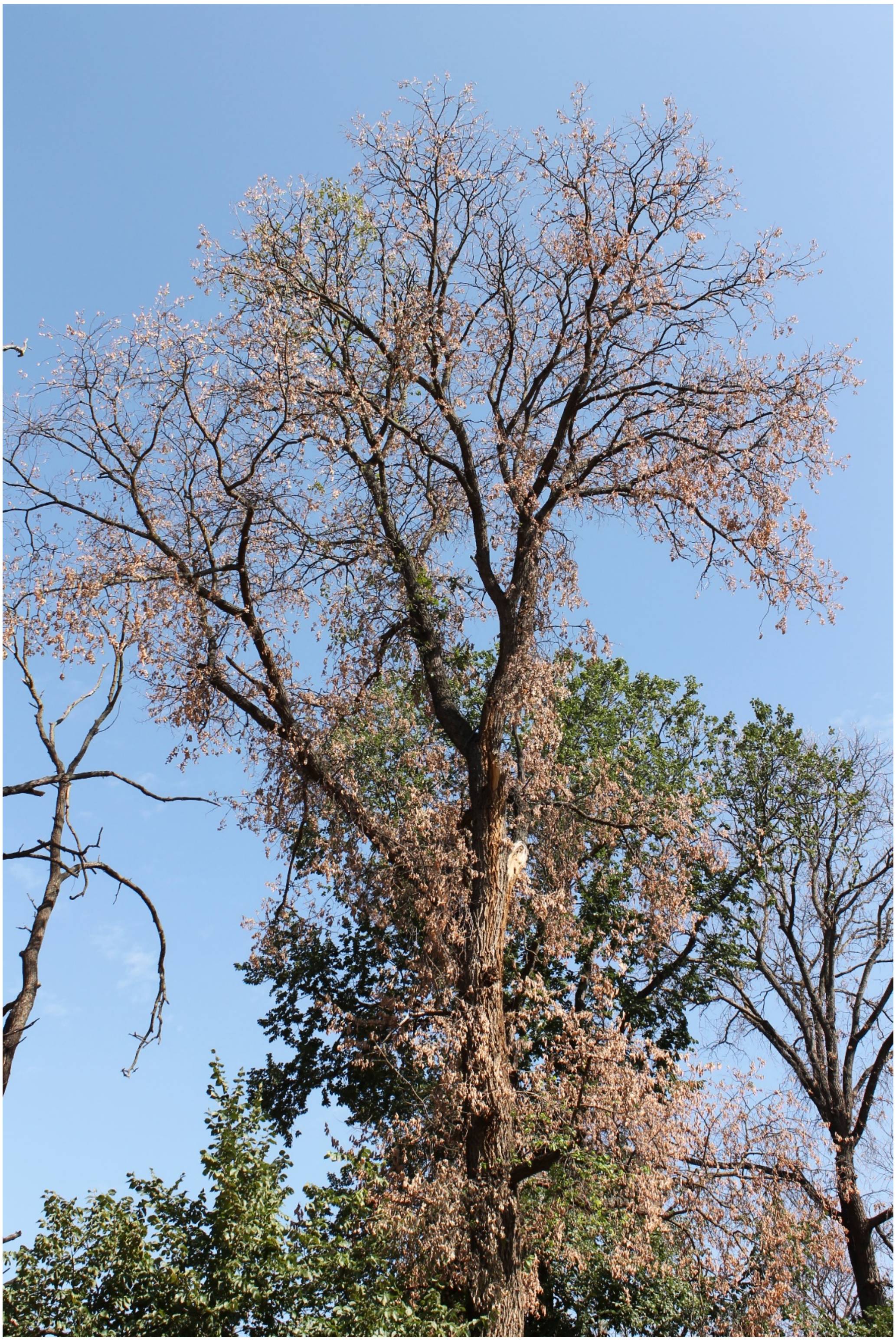

C) RIV3

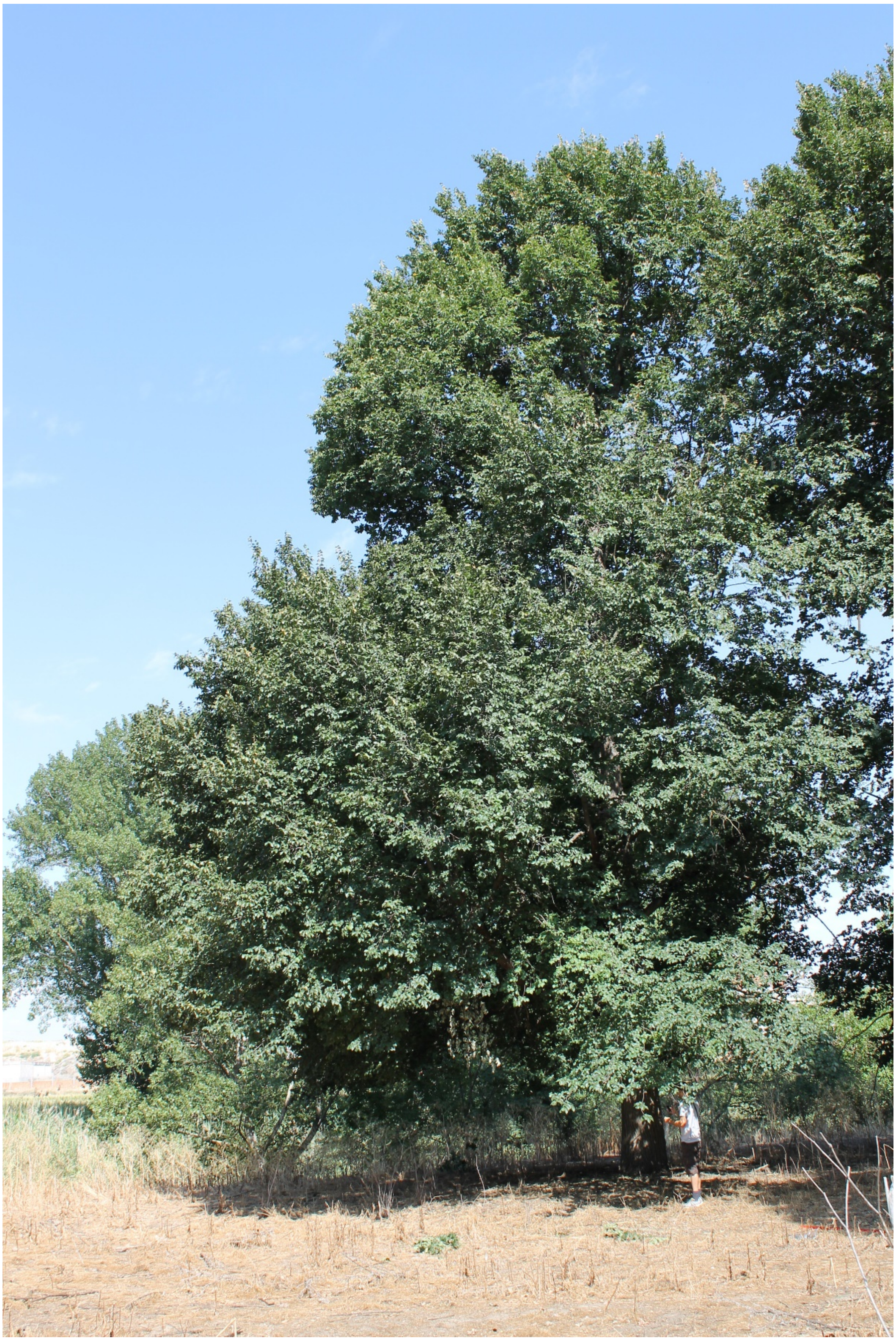

D) RIV4

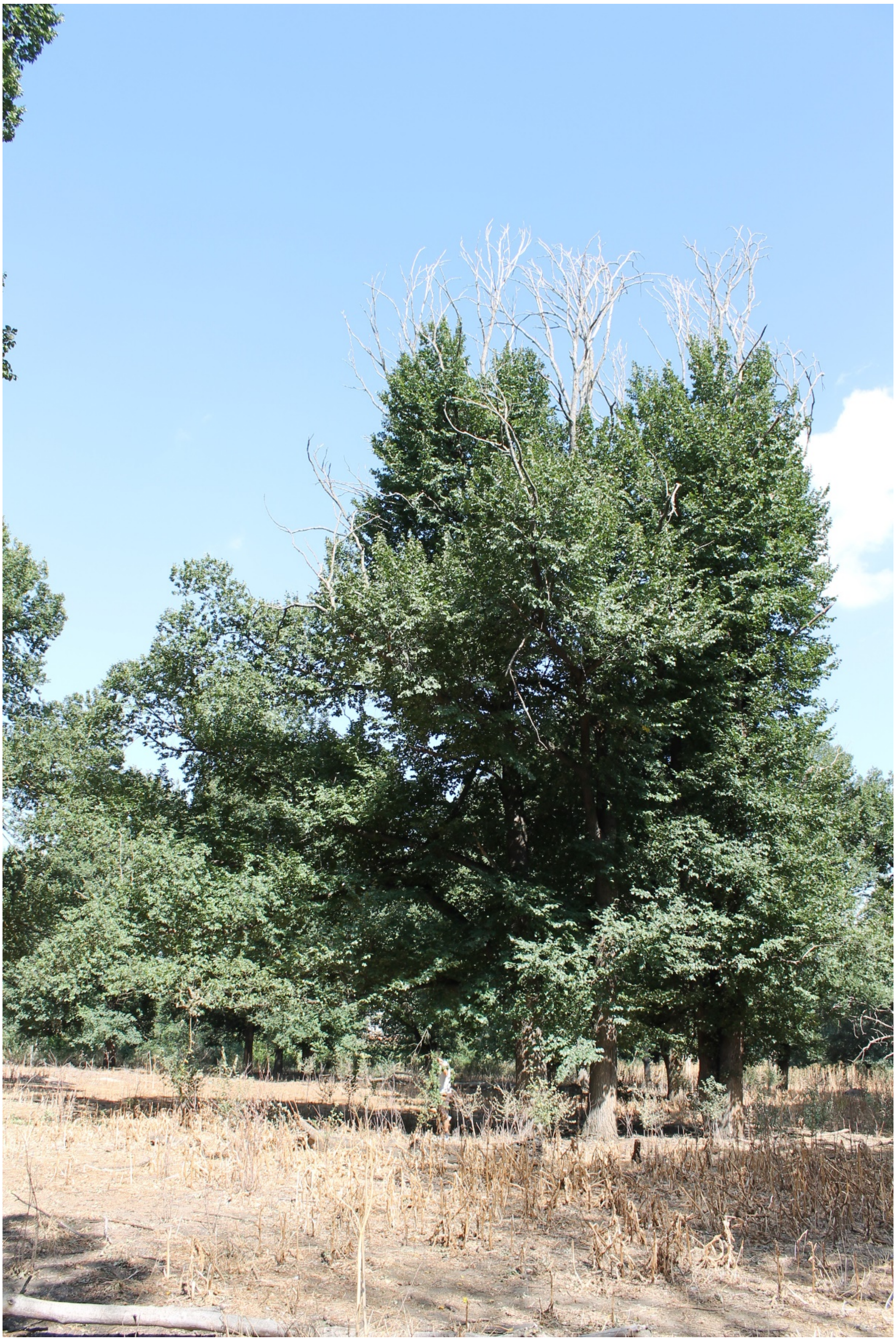

E) RIV5

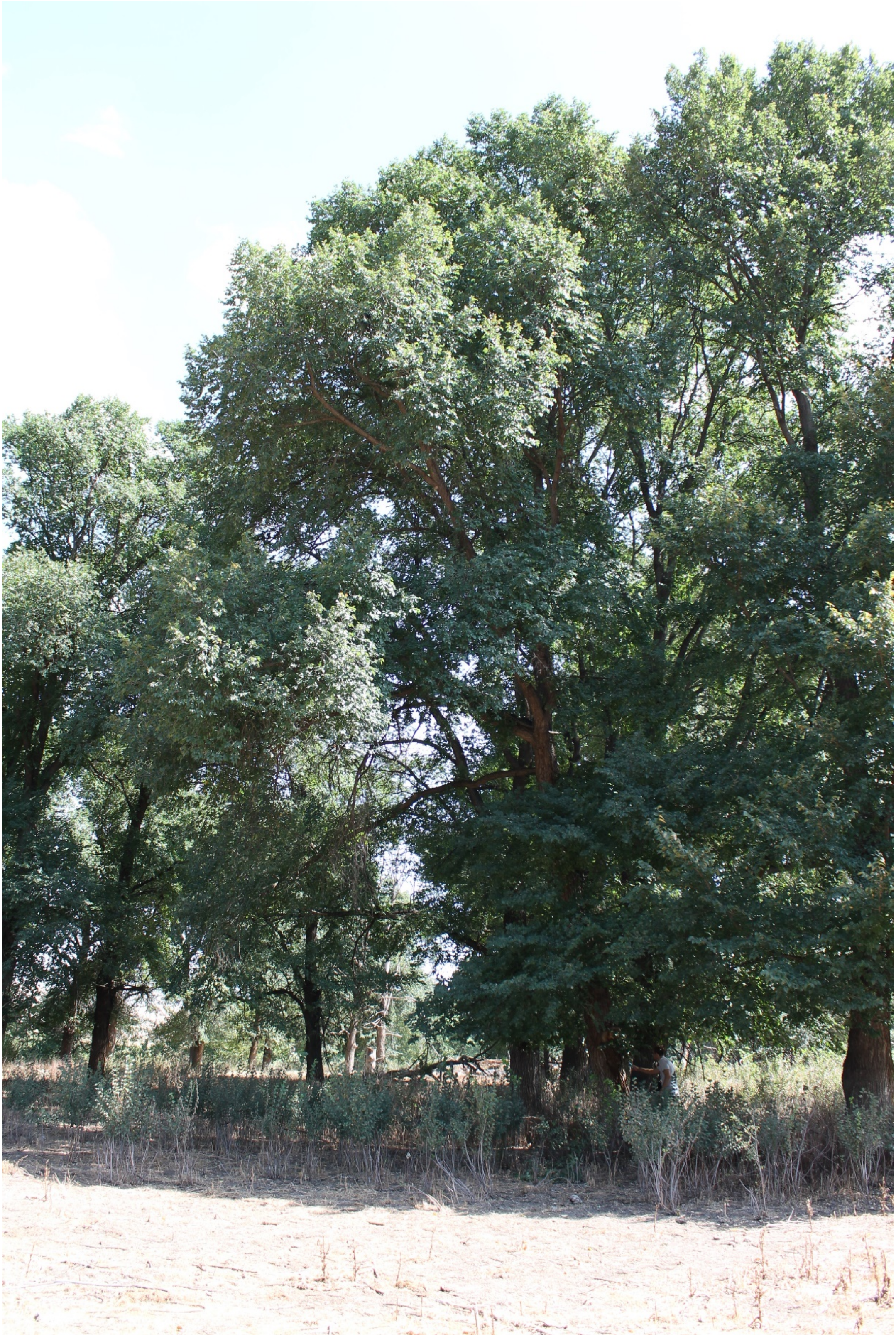

F) RIV6

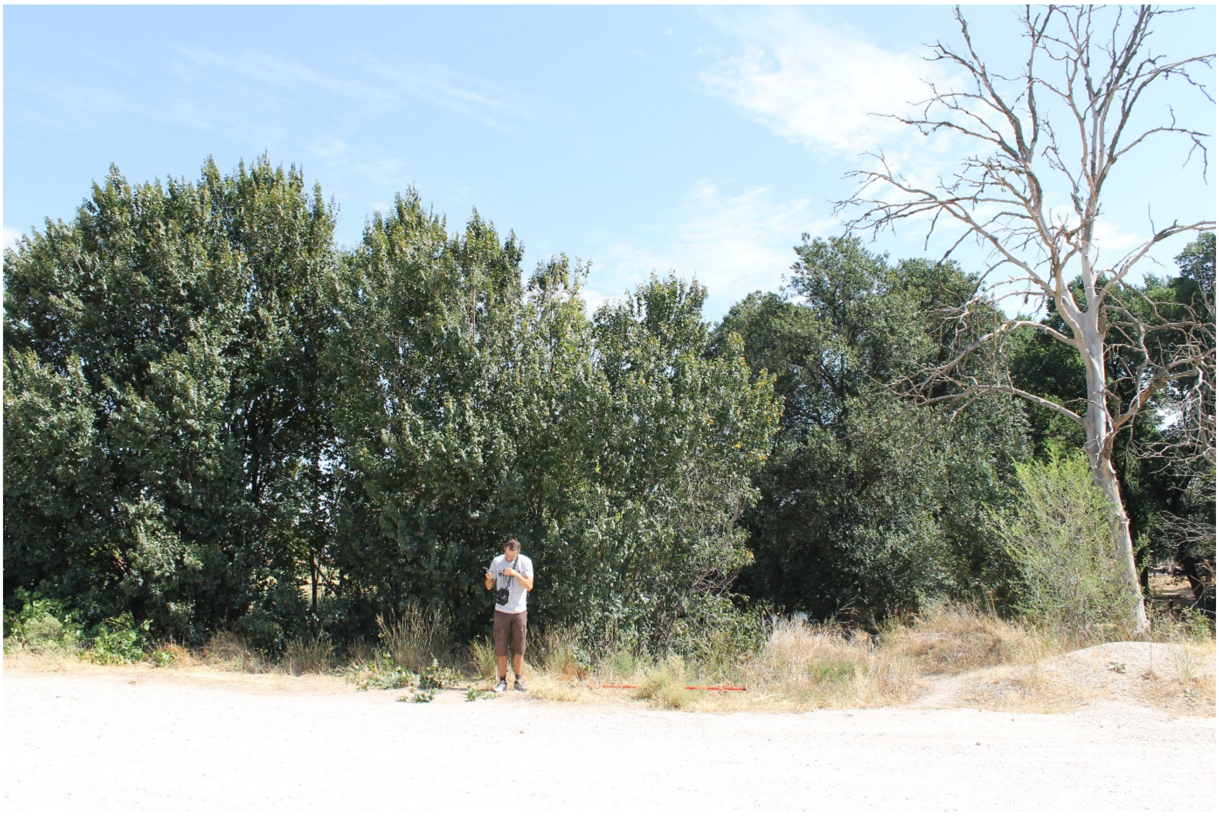

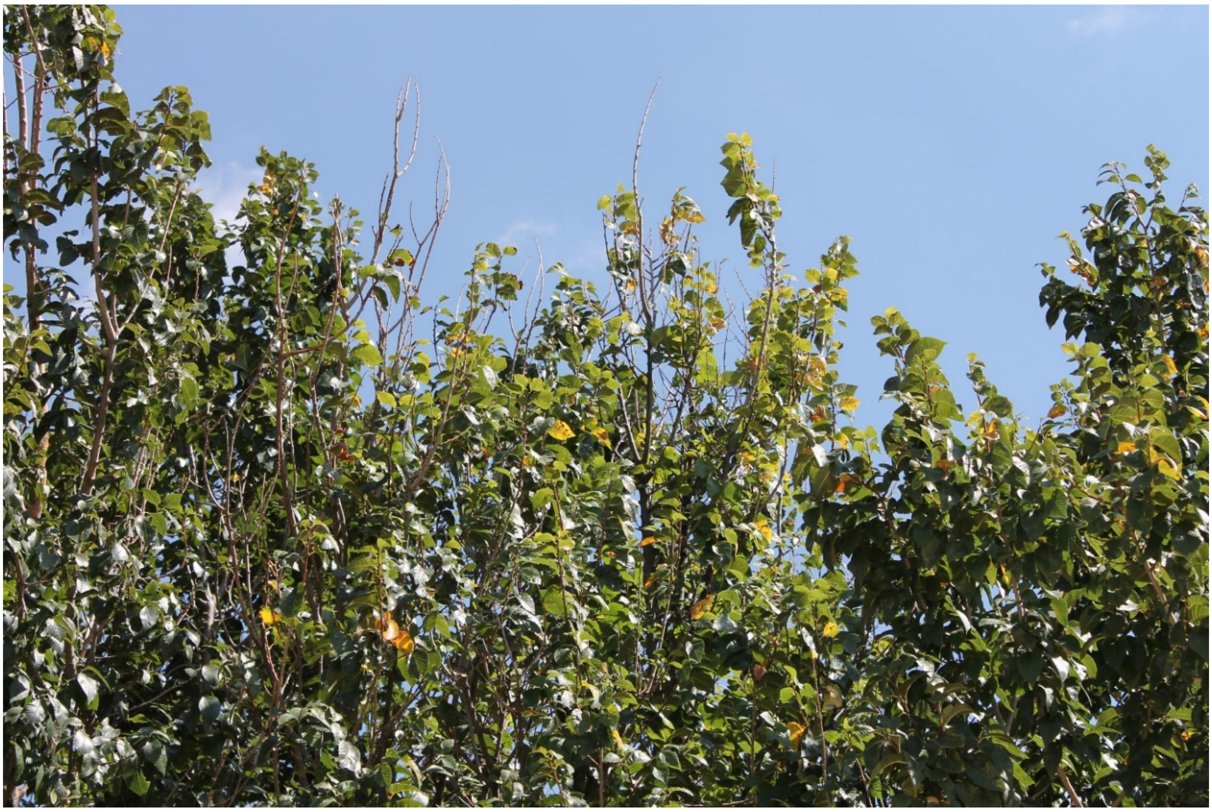

**Sup. Fig. 1.** Trees from the Rivas natural population during sample collection: A) RIV1; B) RIV2; C) RIV3; D) RIV4; E) RIV5; F) RIV6.

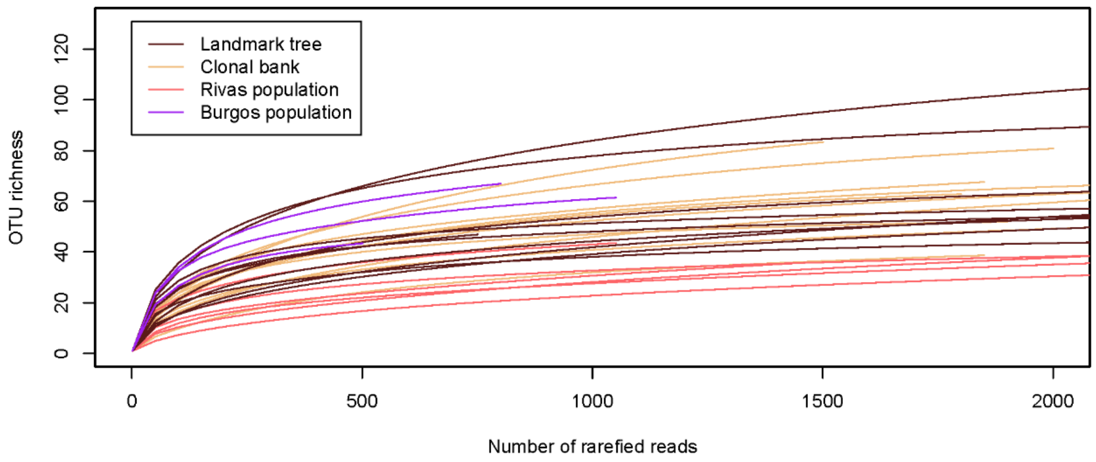


**Sup. Fig. 2.** Rarefaction curves for the 29 samples collected, displaying an increasing sequencing effort from 0 to 2000 reads.

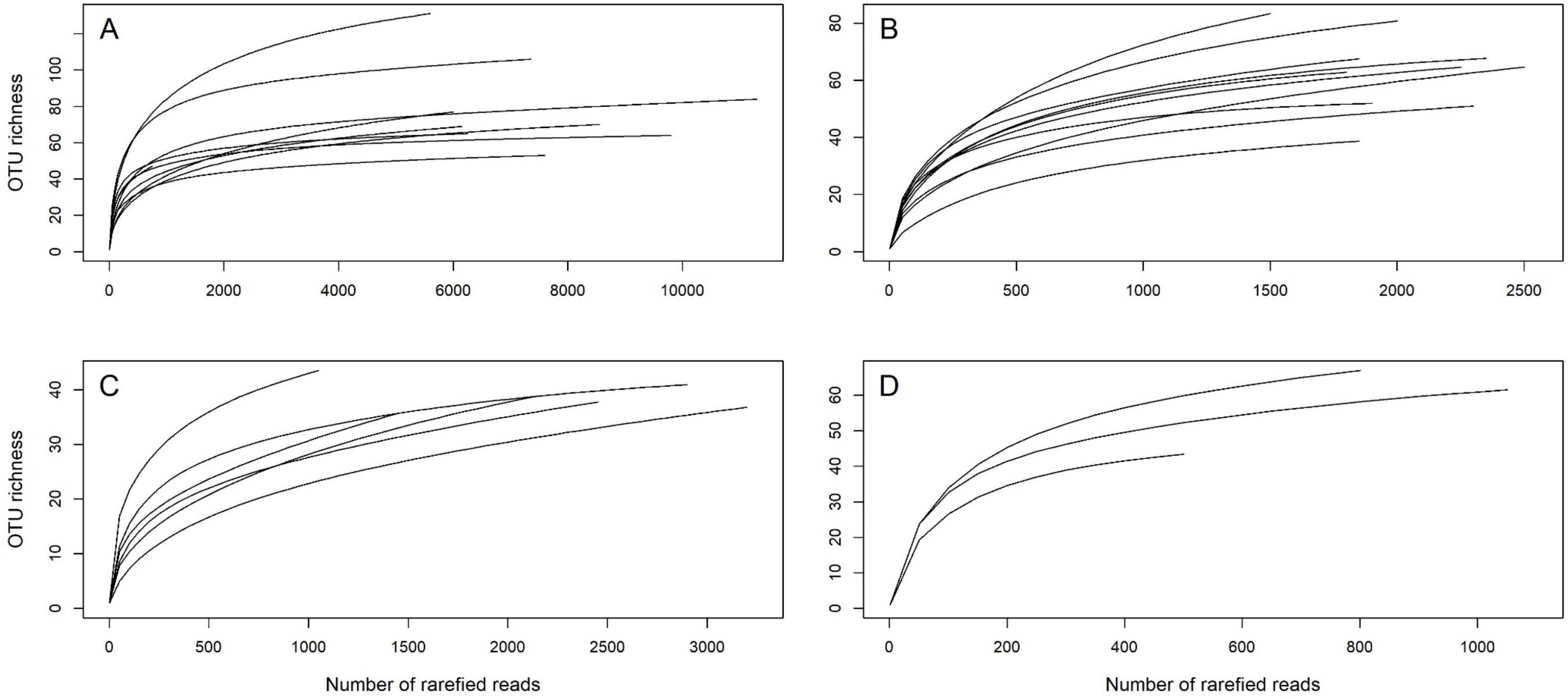


**Sup. Fig. 3.** Rarefaction curves for the samples collected, grouped by collection site, showing an increasing sequencing effort. A) Landmark tree; B) Clonal bank; C) Rivas population; D) Burgos population.

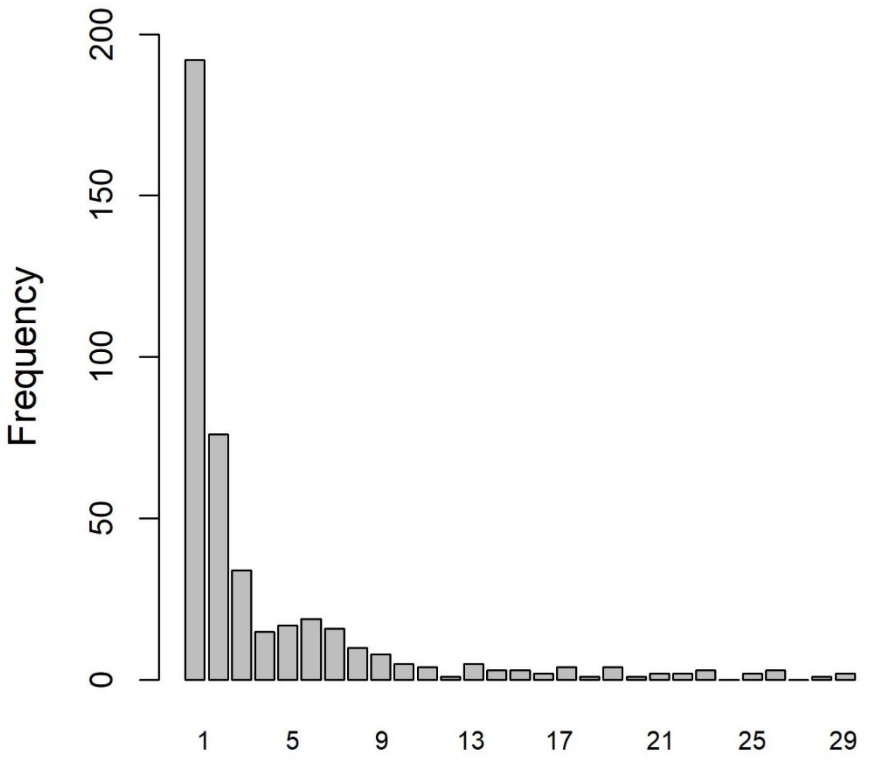


**Sup. Fig. 4.** OTU frequency spectra for total sample set.

**Sup. Table 1.** Susceptibility to DED, measured as a percentage of the crown displaying wilting, shown by the 10 selected *Ulmus minor* genotypes at Puerta de Hierro Forest Breeding Centre.

| Genotype | Origin in Spain | Leaf wilting percentage in screening tests for resistance to DED (and number of replicates) | | | | | | Average |
|---|---|---|---|---|---|---|---|---|
| | | 2002 | 2008 | 2009 | 2010 | 2011 | 2012 | |
| CR-RD2 | Ruidera, Ciudad Real | 93.7 ± 8.3 (8) | - | - | - | - | - | 93.7 ± 8.3 |
| GR-DF3 | Deifontes, Granada | 37.6 ±8.9 (7) | - | 28.7 ± 4.8 (9) | 47.7 ± 7.4 (9) | 34.4 ± 4.4 (9) | - | 37.1 ± 4.0 |
| GR-HL2 | Huélago, Granada | 81.2 ± 8.3 (8) | - | - | - | - | - | 81.2 ± 8.3 |
| J-CA2 | Jaen, Cazorla | 30.0 ± 7.4 (8) | - | 29.0 ± 5.5 (10) | 40.5 ± 6.7 (10) | - | - | 33.2 ± 3.7 |
| MA-PD2 | Málaga, Pedrizas | 36.2 ± 8.3 (8) | - | - | - | 43.5 ± 4.1 (10) | 49.5 ± 7.8 (10) | 43.1 ± 3.8 |
| M-DV1 | Dehesa de la Villa, Madrid | 82.5 ± 8.4 (7) | 89.2 ± 6.4 (10) | 87.1 ± 5.4 (10) | - | 86.7 ± 4.9 (8) | 80.0 ± 6.3 (8) | 85.1 ± 1.7 |
| M-DV5 | Dehesa de la Villa, Madrid | 28.1 ± 8.3 (8) | - | 12.5 ± 5.6 (10) | 13.5 ± 6.9 (10) | - | - | 18.0 ± 5.0 |
| M-RT1.5 | Retiro, Madrid | - | 13.2 ± 2.6 (3) | 19.8 ± 7.9 (3) | - | 14.0 ± 5.2 (7) | 14.2 ± 5.8 (7) | 15.3 ± 1.5 |
| TO-PB1 | Toledo, Puebla de Montalbán | - | - | - | - | 64.5 ± 5.4 (6) | 100.0 ± 0.0 (6) | 82.2 ± 17.8 |
| V-AD2 | Ademuz, Valencia | - | 10.9 ± 7.0 (10) | 17.2 ± 4.6 (10) | - | - | - | 14.0 ± 3.2 |

**Sup. Text 1.** Detailed methods

*Level of tolerance of Ulmus minor genotypes*

Several replicates of each clone (3-10) planted in field experimental plots were artificially inoculated with an aqueous suspension of O. novo-ulmi blastospores (106 spores ml-1), following the methodology described in Martin et al. (2015). All screening tests were performed with trees of at least four years of age. The average percentage of wilting leaves in the crown recorded at day 60 after inoculation was used as a proxy to the tolerance to DED of each clone.

*DNA isolation*

All the samples were stored below 10°C and processed in a laminar flow chamber within the next 24 hours. First, they were surface-sterilized, with sequential baths: (1) ten minutes lightly shaken in autoclaved distilled water with a drop of Tween 20, (2) once dried, 30 seconds in ethanol 75%, (3) five minutes in sodium hypochlorite 4%, and (4) 15 seconds in ethanol 75% and left to dry. Afterwards, they were carefully peeled with a sterilized scalpel, removing the periderm. The wood samples were wrapped in autoclaved aluminum foil and stored at –80°C. Later, the samples were ground with an IKA A11 mill (Staufen, Germany), keeping them frozen with the regular addition of liquid nitrogen. The wood powder was stored at –80°C in sterile centrifuge tubes. The two wood cores from the trunk of the landmark tree had the rhytidome removed. Later they were ground with autoclaved pestles and mortars with regular addition of liquid nitrogen to keep the tissue continuously frozen.

DNA isolation was done after wood powder enzymatic digestion to improve recovery of fungal DNA. Approximately 150 mg of wood powder was incubated for 20 minutes at 37°C in 400 µl of enzymatic solution containing 0.04 mg of Lyticase (ref. n. L2524; Sigma-Aldrich, St. Louis MO, USA) and Tris-EDTA 50 mM. Afterwards, the samples were centrifuged at 13,000 rpm and the supernatant removed. The rest of the DNA isolation was carried out using the Invisorb Spin Plant Mini Kit (STRATEC Biomedical AG, Birkenfeld, Germany) following manufacturer instructions, with the following modifications: after Lysis Buffer, a pinch of zirconium oxide beads 0.1 mm diameter (ref. n. 1005-02; Bertin Instruments, France) was added, and during the subsequent 65°C bath the samples were vortexed three times. These modifications were done to increase fungal cell wall disruption, therefore improving the recovery of fungal DNA.

*DNA amplification*

The profiling of the endophyte composition was carried by high throughput sequencing of the first internal transcribed spacer region (ITS1) of the ribosomal DNA. The amplification was done in two steps. First, we amplified the target region with oligonucleotides that contained the specific fungal primer ITS1-F (Gardes and Bruns 1993) or the non-specific primer ITS2 (White et al. 1990) on the 3' edge and universal tails on the 5' edge. Afterwards, the amplified product was re-amplified with oligonucleotides containing such universal tails, a sample-specific tag and the adaptors for 454-pyrosequencing chemistry. Given that amplifications of the two main sample sets (monumental tree and clonal bank) were done at different times, we were forced to use different polymerases (TaKaRa PrimeSTAR GXL and Roche FastStart High Fidelity, respectively), so the protocols varied slightly. This could rise some concerning in terms of comparative power. However, since this study just pretended to contrast these two datasets primarily in terms of incidence, and not of abundance, different PCR chemistries do not pose a problem. The populations of Rivas-Vaciamadrid and Burgos were assayed with the same conditions as the ones of the clonal bank. Such conditions for the first and the second PCR respectively were: a starting denaturation at 95°C for 2 min; then 35 cycles of 95°C for 30 s, 55°C for 30 s and 72°C for 1 min; finishing with a final elongation step of 72°C for 5 min. For both PCRs, the reactions were done for a volume of 15 µL with final concentrations of ~5ng/µL of total isolated DNA or ~0.1ng/µL of product from the first PCR (1 µL of stock DNA solution or 1 µL of product of the first PCR diluted 1/100), 1× of reaction buffer, 1.8 mM of $MgCl_2$, 0.2 mM of each dNTP, 0.4 mM of each primer and 0.05 U/µL of FastStart enzyme. The conditions for the PrimeSTAR polymerase were as specified in the manufacturer manual for the 3-step PCR, setting Tm to 58°C in the first PCR and 60°C in the second, and increasing the cycles to 35 in the first PCR. Reaction volume was 15 µL.

*High-thoughput sequencing*

We decided the survey on the landmark tree to be the first one because depicting the endophytic mycobiome within a tree would provide information about the sampling effort needed in subsequent experiments to properly characterize each tree mycobiome. Since we did not have a priori information about how deep we should sample, and we considered that deciphering the within-tree endophytic structure deserved a higher resolution, we allotted a large sequencing capacity to this purpose. Given that in the clonal bank experiment we aimed at detecting possible associations to DED tolerance not only of the abundant endophytes but also of the rare ones, such experiment also received an important sequencing allocation to attain a considerable resolution. The elm population of Burgos, on the other hand, was shallowly sequenced, because it was used mostly as an outgroup, to reveal how ubiquitous the core microbiome detected in the other three populations was.After the second PCR, the product of all the samples was quantified, pooled equimolarly and pyrosequenced in a 454 GS FLX Titanium platform (Roche, Switzerland, Basel). The sequencing effort was as follows: monumental tree samples were run in 1/8$^{th}$ of a plate, clonal bank ones in the 62.5% of 1/8$^{th}$ of a plate, Rivas-Vaciamandrid 37.5% of 1/8$^{th}$ of a plate and the Burgos population was run in the 4% of 1/4$^{th}$ plate.

*Default values for* RunTitanium *script from AmpliconNoise v1.29 (Quince et al. 2011)*

Parameters spyro=60 and cpyro=0.01 for PyroNoise; sseq=25 and cseq=0.08 for SeqNoise; and alpha=-7.5 and beta=0.5 for Perseus. The minimum and maximum size of flowgram sizes were 360 and 720 bp, respectively. Sequences were truncated to 400 bp.

*First step in the taxonomic assignment: compare to public database.*

Firstly, we compared the ITS1 sequences of each OTU to the publicly available, curated database of taxonomic information of ITS barcodes UNITE version 7.2 (released at 1-Dec-2017; Kõljalg et al. 2005; Nilsson et al. 2018) using MegaBLAST algorithm (Zhang et al. 2000). The output was parsed using a customized R script. The taxonomic rank of each OTU was given using the best hit but only if the hit covered more than 90% of the query sequence and the similarity was above 97% for species assignment (Blaalid et al. 2013), 95% for genus, 93% for family, 90% for order, 85% for class and 75% for phylum. Otherwise the taxonomic rank was deemed unknown. These varying similarity thresholds were adopted to adjust the level of stringency to the taxonomic rank. Setting a high threshold for all would produce a high level of accuracy in all the ranks, but an unjustified number of missing values in higher ranks as phylum. In contrast, setting an overall low threshold would assign full taxonomic ranks to most OTUs, but lower ranks would be mostly inaccurately assigned. Note that this approach is conservative, since the assignment is always referred to the highest hit. These thresholds were used to avoid assigning taxonomy when only low similarity hits occurred. For that reason, except for species (Blaalid et al. 2013), the rest of thresholds were arbitrary and conservatively chosen. Singletons were not included in these analyses, since they would have provided considerable noise.

*Second step in the taxonomic assignment: haplotype identity of well conserved regions.*

For each OTU we selected an exemplary sequence and retrieved its whole original read (i.e. before the stripping out step performed by FungalITSExtractor). We did that to recover the small portions of 18S and 5.8S ribosomal subunits amplified by the ITS1-F and ITS2 primer pair. These portions of the ribosomal subunits are well-conserved and can introduce bias during the clustering step to form the OTUs (Nilsson et al. 2010). However, for the same reason they can provide very helpful information when assigning OTUs to higher taxonomic ranks. Their slow molecular evolution will produce few, but highly private unique event polymorphisms and, thus, highly discriminant for some taxonomic groups. Having then a complete reference read per OTU, we first inferred the phylum. We aligned the whole set of reference reads using MUSCLE (Edgar 2004) with default settings implemented in MEGA-X v. 10.0.5 (Kumar et al. 2018). Later, we extracted the conserved regions using Gblocks v0.91 (Castresana 2000; Talavera and Castresana 2007) setting the maximum number of contiguous non-conserved positions to 5, the minimum block length to 8 and allowing all gap positions. The output was processed in R using Biostrings v2.46.0 (Pagès et al. 2017) from Bioconductor. We first kept positions covered by at least 80% of the OTUs, and then selected the most discriminant positions between phylum. To pinpoint them, we used only the information from the OTUs with assigned phylum. For each position, we calculated the standard deviations of frequency of all the four possible nucleotides across all the phyla present in this dataset and added them all, having a score per position that was used as index of discrimination. Positions with monomorphic nucleotides bore low indexes, and position with fixed alternate nucleotides among phyla yielded high values. Haplotypes were created with these highly-discriminant positions and haplotype frequency was estimated in each of the phyla. Haplotypes with frequencies over 0.7 were deemed as sufficiently discriminant and all the OTUs with unknown phylum possessing these discriminant haplotypes were assigned to the respective phylum. After subsetting the OTUs to respective groups, same procedure was applied to infer the unknown classes within Ascomycota and Basidiomycota, and order within Dothideomycetes, Sordariomycetes and Agaricomycetes. Since this method uses the OTUs with assigned taxa as a reference, only the taxa with large representation in our dataset could be inferred.

*Visualization of data and test of hypothesis with DeSeq2*

To visually explore for general patterns and groupings, we used the multivariate methods included in the package. First, we apply to the count data a Variance Stabilizing Transformation. Later, the adjusted counts were used to reduce the dimensionality of the dataset through a Principal Component Analysis and to cluster the samples. Afterwards, we test the hypothesis whether any of the taxonomic groups was associated with the tolerance to DED. We implemented the model in DESeq2 setting wilting (a proxy of DED tolerance) as the only explanatory variable. These tests were conducted at OTU, family, order and class levels, and only for groups with more than 5 counts adding up the ten clonal bank samples. Significance was calculated with a Wald test and adjusted for Multi-testing using the default DESeq2 approach that estimates False Discovery Rate adjusted P-values.

At the OTU level, we deemed as significant but to a lower degree passing the threshold of FDR < 0.15: though being significant in the independent tests, the significance was weaker when accounting for multiple-testing. We consider those reports with recording.

**Sup. Text 2.** Supplementary results on sequencing effort, taxonomic assignment and taxonomical composition

*Sequencing effort by sample set*

Sampling coverage was larger for the Somontes tree, accounting for 65.59% of the total reads, reflecting the greater sequencing effort for these samples. The clonal bank collection accounted for 19.44% and the samples from the Rivas population accounted for 12.77%. The three samples from the distant population of Burgos province totalled 2.30% of reads, as expected by the experimental design. One of these samples obtained only 541 reads.

*Improvements in taxonomic information screening highly conserved runs of the ITS1 and flanking sequences*

We were able to infer the higher taxonomic ranks of a considerable number of OTUs. Starting from the top of the taxonomic hierarchy, we could infer the identity of 75 OTUs out of the 83 whose phyla were unidentified only by BLAST on the UNITE database. Those accounted for 18,777 reads (17.7 % of the total number of reads). Regarding class, the next taxonomic rank, we could assign identity to 23 out of 106 unidentified OTUs (2,744 reads, 2.6% of the total number). Finally, we infer the order of 23 out of 157 unidentified OTUs (2,366 reads, 2.2% of the total number). Albeit the gains in the class and order ranks were marginal, this methodology allowed us to determine the phylum of almost all our dataset.

*Taxonomy of the landmark tree endomycobiome*

We were able to assign the taxonomic family to 141 OTUs (59.18% of the reads; 76 families), the order to 192 OTUs (74.41% of the reads; 38 orders), the class to 229 OTUs (80.22% of the reads; 17 classes) and the phylum to 282 OTUs (99.99% of the reads; four phyla).

The phyla included Ascomycota and Basidiomycota, but also Mortierellomycota and Mucoromycota (Fig. 4). The last one was only present in the sample from the north side of the trunk, and Mortierellomycota was only present there and in the close lowest northern branch. Ascomycota accounted for 192 OTUs (87.87% of the taxonomically assigned reads), and Basidiomycota for 87 OTUs (12.00% of the assigned reads). The samples collected from the lowest branches had the highest share of Basidiomycota.

Among Ascomycota, the most common classes were, by order of abundance, Dothideomycetes, Eurotiomycetes, Sordariomycetes, Leotiomycetes, and Lecanoromycetes (Fig. 4). Interestingly, 34 OTUs identified as Ascomycota did not have assignable class. The Basidiomycota were primarily represented by Tremellomycetes, Agaricomycetes, and Cystobasidiomycetes. Twenty-three of the Basidiomycota OTUs had not identified class. All but one of these eight classes including Ascomycota and Basidiomycota were present in all the samples. Agaricomycetes was not found in three spots (the trunk samples and the southern lowest branch).

The most common orders, accounting each for at least 5% of the assigned reads, were Pleosporales, Chaetothyriales, Xylariales, Helotiales, and Capnodiales, all of them Ascomycota. The most common Basidiomycota order was Tremematales accounting for 4.79% of the reads, followed by Filobasidiales (3.22% of the reads). Finally, Herpotrichiellaceae, Didymellaceae, Pleosporaceae, Diatrypaceae, and Cladosporiaceae were, by order, the most common families (each with at least 5% of the assigned reads), all Ascomycota as well. Diatrypaceae was only represented by one single OTU present abundantly in one sample (a middle crown branch) and a singleton in another one. The within-tree pervasive OTUs belong mostly to these just mentioned taxa, albeit ten (out of twenty-three) had unassigned families (Table 2).

*Taxonomy of the clonal bank endomycobiome*

Using UNITE database, combined with our haplotype-based inference, we could assign the taxonomic family to 111 out of 213 OTUs (56.45% of the reads; 61 families), the order to 147 OTUs (74.48% of the reads; 29 orders), the class to 177 OTUs (78.82% of the reads; 15 classes) and the phylum to 211 OTUs (99.98% of the reads; two phyla).

Solely phyla Ascomycota and Basidiomycota were detected (Fig. 5a). The former was represented by 149 OTUs (90.89% of the taxonomically assigned reads), and the latter by 62 OTUs (9.11% of the assigned reads). No tree had an unusually high share of Basidiomycota reads, but both TO-PB1 and CR-RD2 displayed residual counts (below 2%).

The most frequent Ascomycota classes were the same as in the Somontes tree, though the Eurotiomycetes were less common than the Sordariomycetes and the Leotiomycetes. Dothideomycetes was by far the most common group (Fig. 5a). The Orbiliomycetes were quite abundant, being represented by one single OTU (OTU_13) that was present in eight trees. The pattern in Basidiomycota resembled that found in the Somontes tree, but with much sequences assigned to Cystobasidiomycetes than to Agaricomycetes.

The five most common orders were Pleosporales, Xylariales, Myringiales, Dothideales and Caliciales, consistently Ascomycota. Chaetothyriales order was relatively less common in the clonal bank that in the Somontes tree. The most common Basidiomycota order was conspicuously Tremellales. The most common families were Cucurbitariaceae, Diatrypaceae, Aureobasidiaceae, Didymellaceae and Physciaceae.

*Taxonomy of the core endomycobiome*

In total 32 OTUs passed the criteria for core microbiome membership, 28 of which belonged to Ascomycota, and four to Basidiomycota. Six OTUs had unidentified order (one was Basidiomycota). The Basidiomycota belonged to the order Tremellomycetes (2 OTUs) and Cystobasidiomycetes (1). The Ascomycota were mostly Dothideomycetes (11), but also Eurotiomycetes (4), Sordariomycetes (3), Leotiomycetes (2), Lecanoromycetes (2) and Orbiliomycetes (1). The most common class was the Pleosporales (5), followed by the Capnodiales (3).

**Sup. Text 3.** Taxonomical considerations

*Sequencing effort and level of saturation*

The rarefaction curves revealed that even in the poorly sequenced trees from the Burgos population, OTU saturation effectively occurred (Sup. Fig. 2). Therefore, shallow sequencing (ca. 500 reads) was sufficient to fully characterise the core microbiome. Variation in endophyte composition among samples within the same tree, including locations with very distinctive taxa (Fig. 4), indicated that sampling a single location in a tree to describe its overall endophytic community could introduce bias.

*Main fungal endophyte taxa of U. minor*

This is in line with other studies on forest trees (Arnold et al. 2007; Botella and Diez 2011; Soca-Chafre et al. 2011; Langenfeld et al. 2013; Sanz-Ros et al. 2015). A very similar taxonomic profile was detected in a culture-dependent survey made in Mexican yew (Taxus globosa) comprising a wide variety of plant tissues (Soca-Chafre et al. 2011). In contrast to our work, they detected a few Pezizomycetes (effectively absent in our study) and, comparatively, many more Sordariomycetes. They did not isolate members of the Tremellomycetes and Cystobasidiomycetes. Interestingly, those two groups were not detected in either pines (genus Pinus; Botella and Diez 2011; Qadri et al. 2014; Sanz-Ros et al. 2015) or plum yews (genus Cephalotaxus; Langenfeld et al. 2013), using culture-dependent scrutiny. They were also almost absent in loblolly pine (Pinus taeda) using a mixed approach with isolates and environmental PCR (Arnold et al. 2007). Yews, pines and plum yews are all conifers (gymnosperms). However, Tremellomycetes and Cystobasidiomycetes were present in angiosperm Arctic plants (using pyrosequencing; Zhang and Yao 2015), and in European aspen (genus Populus; by strain isolation; Albrectsen et al. 2018). These basidiomycetous endophytes could be specific to angiosperms or limited in gymnosperms, and interestingly have exhibited association with host tolerance to DED (see below).

At order level, Pleosporales and Xylariales were normally the most common groups of endophytes in other forest trees (Soca-Chafre et al. 2011; Langenfeld et al. 2013; Qadri et al. 2014; Sanz-Ros et al. 2015), in parallel with our findings. Yet, Xylariales was not a very frequent group in European aspen leaves (Albrectsen et al. 2010). At odds with elm and poplar, the order Hypocreales (Sordariomycetes) was found much more commonly in some studies in conifers (Soca-Chafre et al. 2011; Sanz-Ros et al. 2015). In elms, this order was only prevalent in some weakened trees in the Rivas populations (see below).

*Results in Rivas*

The most abundant taxon in RIV3 was OTU_3, a Pleosporal present in all the populations but with a very variable abundance among and within tree (pointing to host colonization through local infections). The most abundant OTU in the quite decayed RIV1 belonged to Diatrypaceae (unknown genus). This OTU was not widespread among the sample set, but when present, usually was very abundant (most common OTU in two trees of the clonal bank but effectively absent in the rest, and in one branch of the landmark tree but absent in the rest).

*Improvements in taxonomic assignments thanks to the in-house method*

One of the main caveats of studies in fungal diversity conducted exclusively by molecular barcoding methods using environmental PCR or high-throughput sequencing is that they rely on public repositories to ascribe the resolved OTUs to taxa (Kõljalg et al. 2005). These repositories are enriched in taxa with immediate societal impact such as human pathogens or crop diseases but provide less support for examination of poorly studied groups like tree endobiomes. The method developed for this study, using the conserved portions of the rDNA and flanking sequences amplified with benchmark primers (ITS1-F and ITS2) to generate discriminant haplotypes to assign the obscure endobiome to higher rank taxa, allowed us to assign phylum to the complete obscure fraction of the clonal bank. This fraction shifted the abundance of Ascomycetes from 92.9% to 90.9%, indicating how neglecting the obscure microbiome can introduce bias in the outcome. Our approach also provided evidence of the occurrence of groups poorly represented in the reference databases. Some haplotypes were detected in abundance in the sampled trees but were not attributable to any taxa due to a lack of representation in the public repository. This issue would create situations in that a somewhat similar taxon to the query OTU is represented in UNITE but the actual target of the OTU is not. Molecular similarity will assign that taxon to the query OTU. Higher levels of taxonomy will be most times right (basing on fundamentals of phylogenetics and molecular evolution), but the lower levels will be wrong in those cases.

*Evidence of the occurrence of groups poorly represented in the reference databases*

None of the three OTUs with haplotype GAGAATAGTAT within Ascomycota were taxonomically assigned using UNITE. If manually BLASTed into GenBank, the hits can be only assigned to uncultured fungi from molecular studies, without taxonomic ascription, hinting at a possible undescribed taxonomic group. The very close haplotype CAGAATAGTAT had eight unidentifiable OTUs using UNITE and twelve identifiable, but they were assigned to four different classes. BLASTed to GenBank, some of them retrieved contentious identities to UNITE. This pattern could also be due to another not previously described group of Ascomycota, because not having a group in the reference datasets may produce inconsistent assignments of the OTUs. An alternative hypothesis could be that this haplotype is ancestral in Ascomycota, and so it is shared between different classes. Another instance was haplotype TTATA within Dothideomycetes. It had none assigned OTU but was present both in Rivas and in Burgos populations (4.2% and 6.3%, respectively). Manually BLASTing its OTUs (OTU_99 and OTU_435) to the website UNITE database v8.0 lead to no in-depth identification but doing so to the GenBank dataset yielded OTU_99 an almost perfect hit in a very recently released sequence (MK442618.1; Crous et al. 2019), identified as a Dictyosporiaceae, a novel ascomycetous family (Boonmee et al. 2016).

**Sup. Text 4.** Supplementary references

Albrectsen BR, Björkén L, Varad A, et al (2010) Endophytic fungi in European aspen (*Populus tremula*) leaves—diversity, detection, and a suggested correlation with herbivory resistance. Fungal Divers 41:17–28. doi: 10.1007/s13225-009-0011-y

Albrectsen BR, Siddique AB, Decker VHG, et al (2018) Both plant genotype and herbivory shape aspen endophyte communities. Oecologia 187:535–545. doi: 10.1007/s00442-018-4097-3

Arnold AE, Henk DA, Eells RL, et al (2007) Diversity and phylogenetic affinities of foliar fungal endophytes in loblolly pine inferred by culturing and environmental PCR. Mycologia 99:185–206

Blaalid R, Kumar S, Nilsson RH, et al (2013) ITS1 versus ITS2 as DNA metabarcodes for fungi. Mol Ecol Resour 13:218–224. doi: 10.1111/1755-0998.12065

Boonmee S, D’souza MJ, Luo Z, et al (2016) Dictyosporiaceae fam. nov. Fungal Divers 80:457–482. doi: 10.1007/s13225-016-0363-z

Botella L, Diez JJ (2011) Phylogenic diversity of fungal endophytes in Spanish stands of Pinus halepensis. Fungal Divers 47:9–18. doi: 10.1007/s13225-010-0061-1

Castresana J (2000) Selection of Conserved Blocks from Multiple Alignments for Their Use in Phylogenetic Analysis. Mol Biol Evol 17:540–552. doi: 10.1093/oxfordjournals.molbev.a026334

Crous PW, Schumacher RK, Akulov A, et al (2019) New and Interesting Fungi. 2. Fungal Syst Evol 3:57–134. doi: doi:10.3114/fuse.2019.03.06

Edgar RC (2004) MUSCLE: multiple sequence alignment with high accuracy and high throughput. Nucleic Acids Res 32:1792–1797. doi: 10.1093/nar/gkh340

Gardes M, Bruns TD (1993) ITS primers with enhanced specificity for basidiomycetes - application to the identification of mycorrhizae and rusts. Mol Ecol 2:113–118. doi: 10.1111/j.1365-294X.1993.tb00005.x

Kõljalg U, Larsson K-H, Abarenkov K, et al (2005) UNITE: a database providing web-based methods for the molecular identification of ectomycorrhizal fungi. New

Phytol 166:1063–1068. doi: 10.1111/j.1469-8137.2005.01376.x

Kumar S, Stecher G, Li M, et al (2018) MEGA X: Molecular Evolutionary Genetics Analysis across Computing Platforms. Mol Biol Evol 35:1547–1549. doi: 10.1093/molbev/msy096

Langenfeld A, Prado S, Nay B, et al (2013) Geographic locality greatly influences fungal endophyte communities in Cephalotaxus harringtonia. Fungal Biol 117:124–136. doi: https://doi.org/10.1016/j.funbio.2012.12.005

Martin JA, Solla A, Venturas M, et al (2015) Seven *Ulmus minor* clones tolerant to *Ophiostoma novo-ulmi* registered as forest reproductive material in Spain. iForest - Biogeosciences For 8:172–180. doi: 10.3832/ifor1224-008

Nilsson RH, Glöckner FO, Saar I, et al (2018) The UNITE database for molecular identification of fungi: handling dark taxa and parallel taxonomic classifications. Nucleic Acids Res 47:D259–D264. doi: 10.1093/nar/gky1022

Nilsson RH, Veldre V, Hartmann M, et al (2010) An open source software package for automated extraction of ITS1 and ITS2 from fungal ITS sequences for use in high-throughput community assays and molecular ecology. Fungal Ecol 3:284–287. doi: https://doi.org/10.1016/j.funeco.2010.05.002

Pagès H, Aboyoun P, Gentleman R, DebRoy S (2017) Biostrings: Efficient manipulation of biological strings

Qadri M, Rajput R, Abdin MZ, et al (2014) Diversity, Molecular Phylogeny, and Bioactive Potential of Fungal Endophytes Associated with the Himalayan Blue Pine (Pinus wallichiana). Microb Ecol 67:877–887. doi: 10.1007/s00248-014-0379-4

Quince C, Lanzen A, Davenport RJ, Turnbaugh PJ (2011) Removing Noise From Pyrosequenced Amplicons. BMC Bioinformatics 12:38. doi: 10.1186/1471-2105-12-38

Sanz-Ros A V, Müller MM, San Martín R, Diez JJ (2015) Fungal endophytic communities on twigs of fast and slow growing Scots pine (Pinus sylvestris L.) in northern Spain. Fungal Biol 119:870–883. doi:

https://doi.org/10.1016/j.funbio.2015.06.008

Soca-Chafre G, Rivera-Orduña FN, Hidalgo-Lara ME, et al (2011) Molecular phylogeny and paclitaxel screening of fungal endophytes from Taxus globosa. Fungal Biol 115:143–156. doi: https://doi.org/10.1016/j.funbio.2010.11.004

Talavera G, Castresana J (2007) Improvement of Phylogenies after Removing Divergent and Ambiguously Aligned Blocks from Protein Sequence Alignments. Syst Biol 56:564–577. doi: 10.1080/10635150701472164

White TJ, Bruns T, Lee S, Taylor J (1990) Amplification and direct sequencing of fungal ribosomal RNA genes for phylogenetics. In: Innis MA, Gelfand DH, Sninsky JJ, White TJ (eds) PCR protocols: a guide to methods and applications. Academic Press, San Diego, pp 315–322

Zhang T, Yao Y-F (2015) Endophytic Fungal Communities Associated with Vascular Plants in the High Arctic Zone Are Highly Diverse and Host-Plant Specific. PLoS One 10:e0130051

Zhang Z, Schwartz S, Wagner L, Miller W (2000) A Greedy Algorithm for Aligning DNA Sequences. J Comput Biol 7:203–214. doi: 10.1089/10665270050081478
